# Canonical and Non-Canonical Psychedelic Drugs Induce Common Network Changes in Human Cortex

**DOI:** 10.1101/2022.10.14.512285

**Authors:** Rui Dai, Tony E. Larkin, Zirui Huang, Vijay Tarnal, Paul Picton, Phillip E. Vlisides, Ellen Janke, Amy McKinney, Anthony G. Hudetz, Richard E. Harris, George A. Mashour

**Affiliations:** Department of Anesthesiology, University of Michigan Medical School, Ann Arbor, MI 48109; Center for Consciousness Science, University of Michigan Medical School, Ann Arbor, MI 48109; Michigan Psychedelic Center, University of Michigan Medical School, Ann Arbor, MI, 48109; Chronic Pain and Fatigue Research Center, University of Michigan Medical School, Ann Arbor, MI, 48109; Neuroscience Graduate Program, University of Michigan, Ann Arbor, MI 48109; Department of Pharmacology, University of Michigan Medical School, Ann Arbor, MI, 48109

## Abstract

The neurobiology of the psychedelic experience is not fully elucidated. Identifying common brain network changes induced by both canonical (i.e., acting at the 5-HT2 receptor) and non-canonical psychedelics would provide mechanistic insight into state-specific characteristics. We analyzed whole-brain functional connectivity based on resting-state fMRI data in humans, acquired before and during the administration of nitrous oxide, ketamine, and lysergic acid diethylamide. We report that, despite distinct molecular mechanisms and modes of delivery, all three psychedelics reduced within-network functional connectivity and enhanced between-network functional connectivity. More specifically, all drugs tested increased connectivity between right temporoparietal junction and bilateral intraparietal sulcus as well as between precuneus and left intraparietal sulcus. These regions fall within the posterior cortical “hot zone,” posited to mediate the content of consciousness. Thus, both canonical and non-canonical psychedelics modulate networks within an area of known relevance for conscious experience, identifying a biologically plausible candidate for their subjective effects.

## Introduction

The neurobiological basis of the psychedelic experience remains incompletely understood. One approach to deeper mechanistic insight would be the identification of drug-invariant neural correlates induced by diverse psychedelic drugs. Canonical or classical psychedelics such as lysergic acid diethylamide (LSD) are thought to exert their effects primarily through the serotonergic 5-HT_2_ receptor, whereas the non-canonical psychedelic ketamine—sometimes referred to as a dissociative drug—acts through glutamatergic NMDA receptors (*1*).

Nitrous oxide is another NMDA receptor antagonist (*2*) that has been in continuous clinical use as an anesthetic since the mid-19^th^ century and that has psychedelic properties at subanesthetic concentrations (*3*). Unlike LSD and ketamine, there is a paucity of data on the neural correlates of the psychedelic experience induced by nitrous oxide, despite longstanding use of this inhaled drug and a description of its psychological effects by William James more than a century ago (*4*). Various electroencephalographic and magnetoencephalographic studies in humans have reported spectral, functional connectivity, and complexity changes associated with nitrous oxide (*5*–*10*), but at sedative rather than psychedelic concentrations or without assessment of psychedelic phenomenology. Although there has been investigation of the effect of nitrous oxide on cerebral blood flow using magnetic resonance imaging (MRI) (*11*), there have been no functional MRI (fMRI) studies during nitrous oxide exposure in humans that have characterized changes in functional brain networks associated with psychedelic effects. Thus, the neural correlates of the psychedelic experience induced by nitrous oxide, and the relationship of such correlates to the neurobiology of other psychoactive drugs such as LSD or ketamine, is unclear.

We conducted a neuroimaging study of healthy human volunteers, who were assessed with a validated altered states questionnaire before and after exposure to psychedelic concentrations of nitrous oxide. We analyzed whole-brain functional connectivity based on MRI data acquired before and during the administration of nitrous oxide. To compare the neural correlates of the psychedelic experience to other drugs, we conducted a secondary analysis of fMRI data acquired during exposure to subanesthetic ketamine and LSD. To control for non-specific pharmacological perturbations of brain networks, we also assessed functional connectivity changes during propofol sedation, which does not evoke psychedelic experiences. We report that, despite distinct molecular mechanisms and modes of delivery, nitrous oxide, ketamine, and LSD all reduce within-network functional connectivity and enhance between-network functional connectivity. More specifically, after excluding network changes induced by propofol sedation, these canonical and non-canonical psychedelic drugs consistently increased connectivity between temporoparietal junction (TPJ) and intraparietal sulcus (IPS), two regions located in the so-called “posterior cortical hot zone” that is thought to mediate content of consciousness. These data support the hypothesis that there is a common, drug-invariant neurobiology to the psychedelic experience.

## Results

### Nitrous Oxide as a Psychedelic

Psychedelic experiences induced by LSD and ketamine have already been well described (*12*, *13*). To characterize the altered state of consciousness induced by 35% nitrous oxide, the 11-D altered states of consciousness questionnaire was performed. Nitrous oxide induced a significant change in each dimension when comparing study score to the pre-nitrous-oxide baseline score: experiences of unity (t (12) = 3.315, FDR-corrected p=0.013), spiritual experience (t (12) = 2.855, FDR-corrected p=0.017), blissful state ((t (12) = 3.692, FDR-corrected p=0.011), insightfulness ((t (12) = 3.487, FDR-corrected p=0.011), disembodiment ((t (12) = 5.302, FDR-corrected p=0.002), impaired control and cognition ((t (12) = 3.066, FDR-corrected p=0.015), anxiety ((t (12) = 2.237, FDR-corrected p=0.045), complex imagery ((t (12) = 3.800, FDR-corrected p=0.011), elementary imagery ((t (12) = 2.375, FDR-corrected p=0.039), audiovisual synesthesia ((t (12) = 3.168, FDR-corrected p=0.015), and changed meaning of percepts ((t (12) = 3.017, FDR-corrected p=0.015). Overall, the altered states of consciousness scores during nitrous oxide administration were higher than pre-nitrous oxide baseline scores on every subscale (Figure 1). Among these 11 dimensions, the change in disembodiment was the most significant, consistent with the designation of nitrous oxide as a dissociative drug.

**Figure 1:**
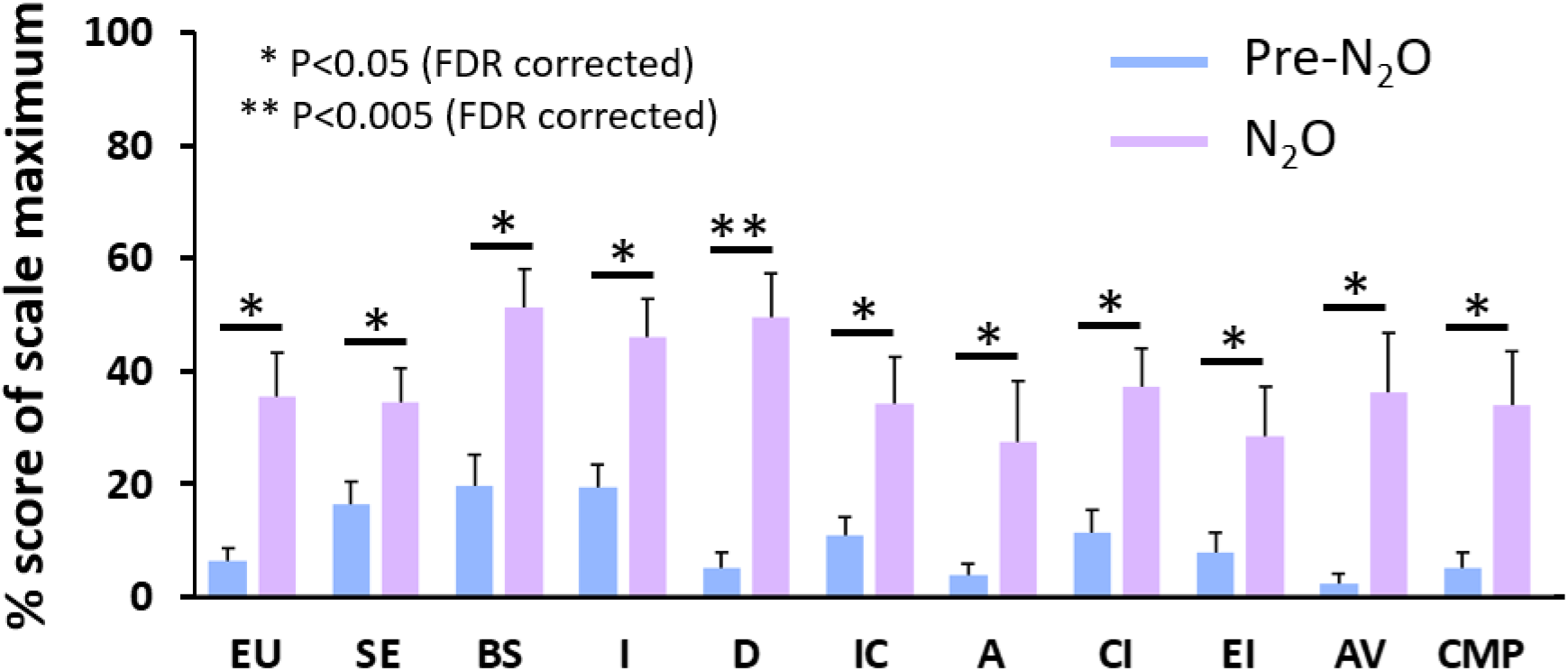
Behavioral results derived from the 11D-Altered States Questionnaire. Error bars represent standard errors. EU: experience of unity, SE: spiritual experience, BS: blissful state, I: insightfulness, D: disembodiment, IC: impaired control and cognition, A: anxiety, CI: complex imagery, EI: elementary imagery, AV: audio-visual synesthesia, CMP: changed meaning of percepts.

### Effects of Nitrous Oxide on Functional Connectivity

Whole-brain ROI-to-ROI functional connectivity during the administration of nitrous oxide was analyzed in comparison with the control condition. Nitrous oxide increased connectivity *between* networks, including visual - salience network (FDR-corrected p= 0.0068), dorsal attention – frontoparietal network (FDR-corrected p= 0.0245), sensorimotor - language network (FDR-corrected p= 0.0245), dorsal attention - language network (FDR-corrected p= 0.0245), salience - default mode network (FDR-corrected p= 0.0245), and dorsal attention – default mode network (FDR-corrected p= 0.0245). Nitrous oxide decreased connectivity *within* salience network (FDR-corrected p= 0.0245) and language network (FDR-corrected p= 0.0245) (Figure 2). For further details, see Table S2.

**Figure 2:**
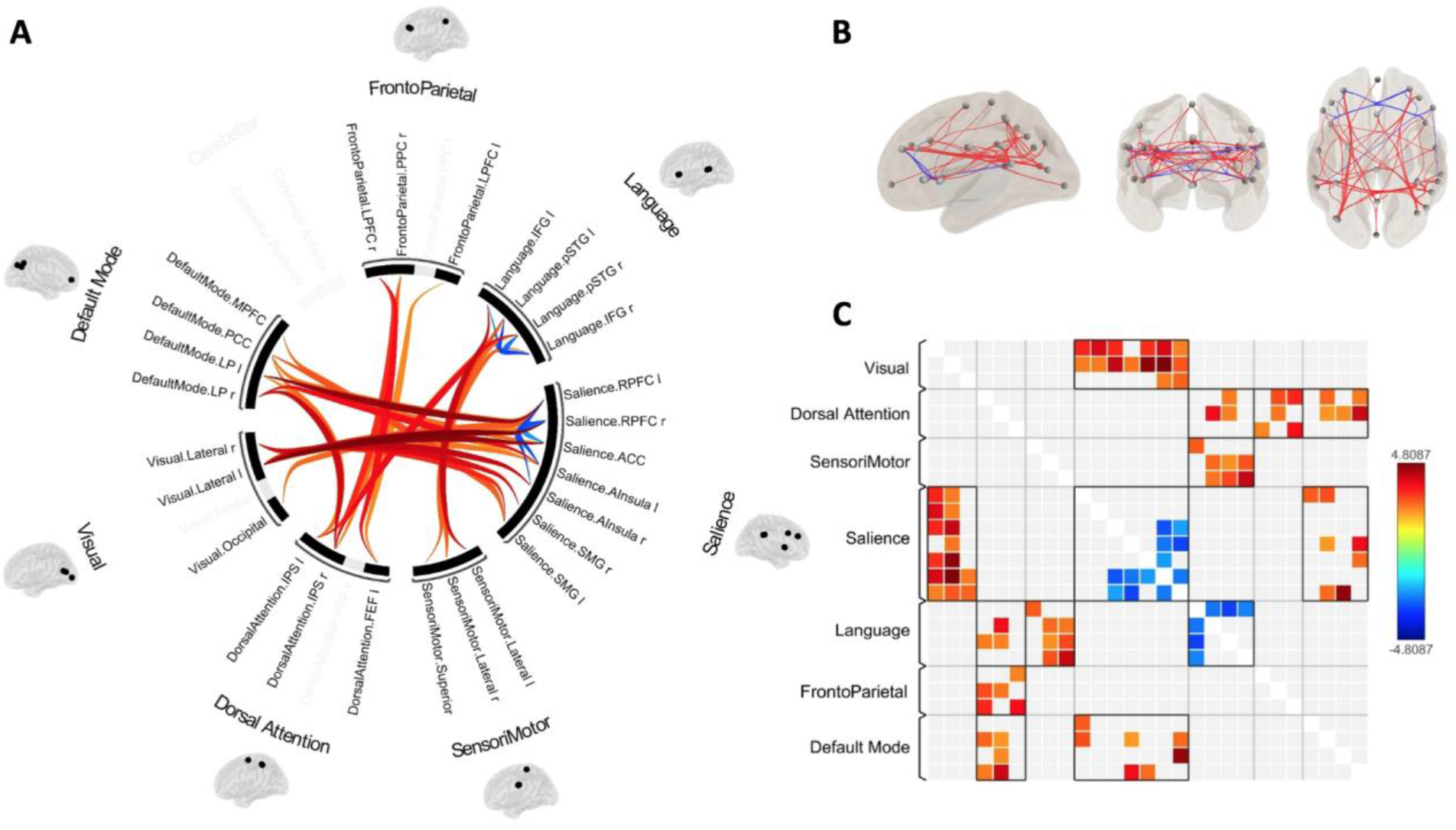
Effects of nitrous oxide on functional connectivity. (A) The circle view displays significant functional connectivity changes (nitrous oxide versus control condition) between

ROIs of seven cerebral cortical networks and one cerebellar network. (B) The connectome view displays the ROIs with individual suprathreshold connectivity lines between them. (C) Depiction of the ROI-to-ROI connectivity matrix of nitrous oxide versus control condition.

### Effects of Ketamine and LSD on Functional Connectivity

To compare the cortical network changes induced by nitrous oxide to those of other non-canonical and canonical psychedelic drugs, we analyzed ROI-to-ROI functional connectivity of the whole brain during exposure to psychedelic doses of ketamine and LSD using a within-group design. Compared to its own baseline, ketamine infusion enhanced between-network connectivity in frontoparietal - default mode network (FDR-corrected p= 0.0198), salience - default mode network (FDR-corrected p= 0.0283), language - default mode network (FDR-corrected p= 0.0306), dorsal attention - default mode network (FDR-corrected p= 0.0446). Ketamine reduced within-network connectivity in frontoparietal network (FDR-corrected p= 0.0198) and sensorimotor network (FDR-corrected p= 0.0273) (Figure 3. A-C and Table S3). Compared to its own baseline (Figure 3. D-E and Table S4), LSD increased between-network connectivity in visual - language network (FDR-corrected p= 0.0030), dorsal attention – language network (FDR-corrected p= 0.0196), language - default mode network (FDR-corrected p= 0.0196), visual - default mode network (FDR-corrected p= 0.0252), dorsal attention – default mode network (FDR-corrected p= 0.0252), salience - default mode network (FDR-corrected p= 0.0350), sensorimotor - default mode network (FDR-corrected p= 0.0423), and frontoparietal - default mode network (FDR-corrected p= 0.0423). LSD decreased within-network connectivity in sensorimotor network (FDR-corrected p= 0.0304) and dorsal attention network (FDR-corrected p= 0.0304).

**Figure 3:**
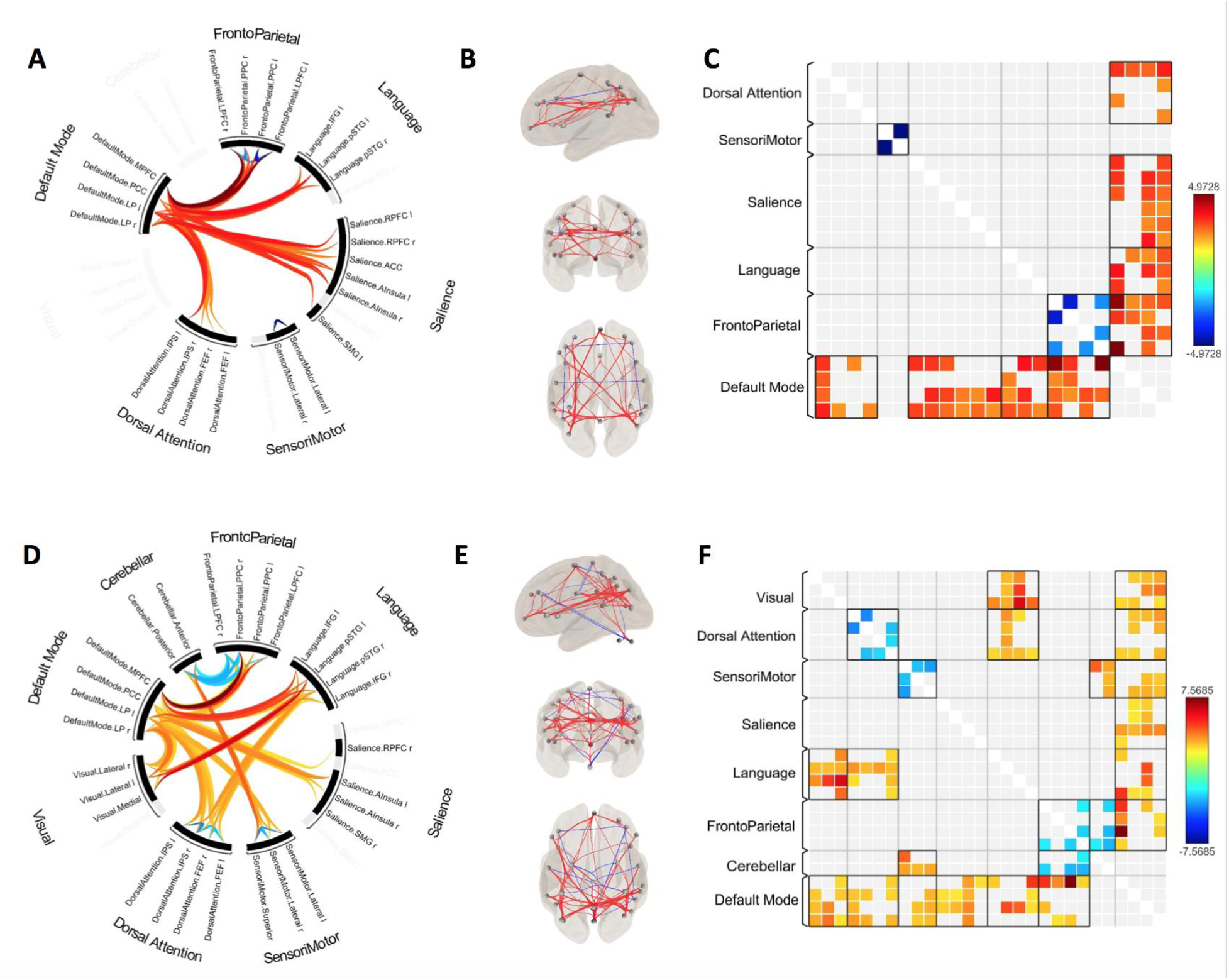
Effects of psychedelic ketamine and LSD on functional connectivity. (A-C) circle view, connectome view, and correlation matrix of functional connectivity changes by ketamine relative to baseline. (D-E) circle view, connectome view, and correlation matrix of functional connectivity changes by LSD relative to baseline.

### Common Effects of Psychedelics on Functional Connectivity

Based on ROI-to-ROI functional connectivity analyses, all three psychedelics decreased within-network connectivity and increased between-network connectivity. However, specific network changes differed across the drugs. Therefore, we assessed whether there were common neural correlates of psychedelic drug administration. Four functional connectivity cluster pairs were consistently affected by all three psychedelics: right lateral parietal/ temporoparietal junction (TPJ) – left intraparietal sulcus (IPS), right lateral parietal – left anterior insula (Ains), right lateral parietal – right intraparietal sulcus, and precuneus cortex – left intraparietal sulcus.

To confirm that these common connectivity patterns were not simply a generic feature of any pharmacologically altered state of consciousness, we analyzed fMRI data during baseline consciousness and propofol sedation as a control condition. Propofol is a clinical anesthetic that, at subanesthetic concentrations, alters consciousness without the typical features of the psychedelic experience. We performed the same whole-brain ROI-to-ROI functional connectivity analysis of the states before and during exposure to propofol sedation. Unlike the psychedelic drugs, there was no evidence of decreased within-network connectivity during subanesthetic propofol administration (Figure S1 and Table S5), and only one functional connectivity cluster pair was consistent with the effect of the three psychedelics: right LP – left Ains (Figure 4A). After eliminating the functional connectivity change also induced by subanesthetic propofol, three common cluster pairs were altered by psychedelic drug administration, including right TPJ/lateral parietal to bilateral IPS and precuneus to left IPS (Figure 4B).

**Figure 4:**
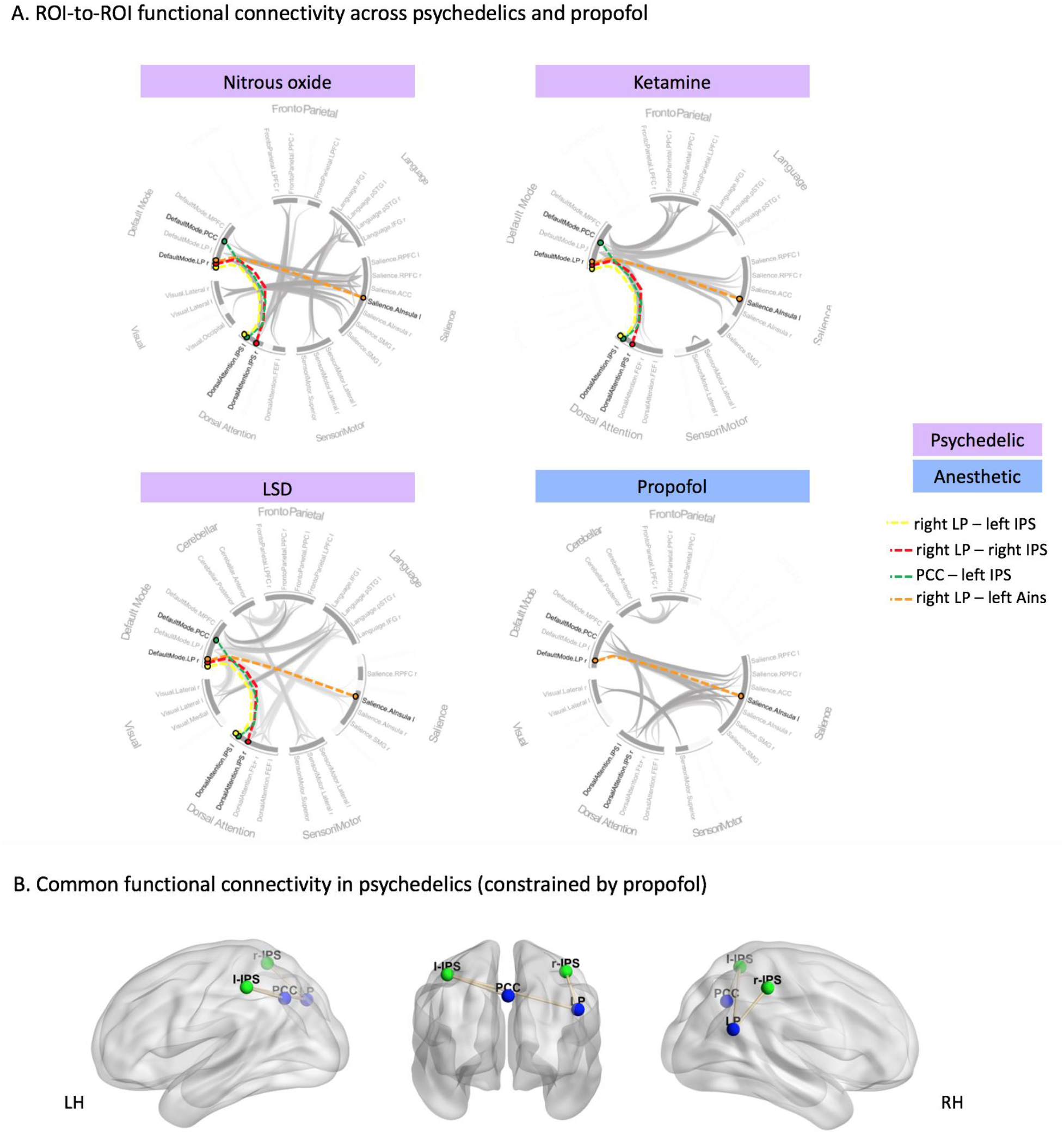
Common effects of psychedelics on functional connectivity. (A) ROI-to-ROI functional connectivity changes induced by nitrous oxide, ketamine, LSD, and propofol. (B) Common functional connectivity patterns due to psychedelic drug administration after removing the change also induced by propofol sedation. LP: lateral parietal cortex, IPS: intraparietal sulcus, PCC: precuneus, Ains: anterior insula, LH: left hemisphere, RH: right hemisphere.

We conducted further analysis of the TPJ because of its association with psychedelic drug administration in this study, and because it is thought to be critical to multisensory integration, consciousness, body ownership, and the psychedelic effects of ketamine (*14*, *15*). We performed a TPJ seed-based functional connectivity analysis in each group and compared the TPJ to the whole brain correlation map between each psychedelic condition and its control condition (Figure 5). The overlap of the TPJ seed-based functional connectivity map across three psychedelics is in the bilateral IPS, which aligns perfectly with the ROI-to-ROI functional connectivity results. In contrast, the TPJ seed-based functional connectivity result of propofol sedation is in the occipital cortex, non-overlapping with the functional connectivity patterns induced by nitrous oxide, ketamine, or LSD.

**Figure 5:**
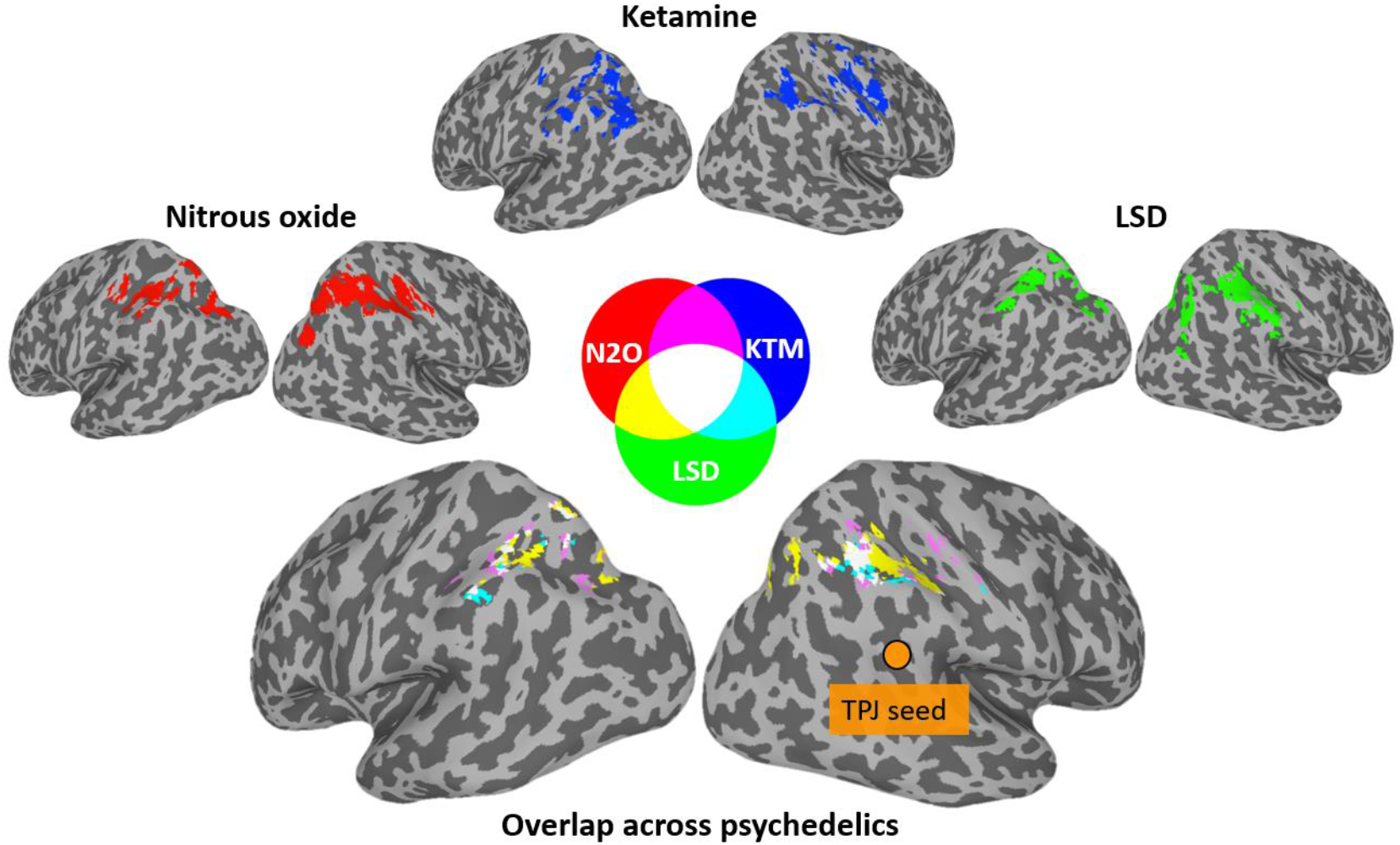
Temporoparietal junction (TPJ) seed-based functional connectivity overlap with nitrous oxide, ketamine and LSD mapped onto an inflated cortical surface. Color code indicates the degree of consistency across the three psychedelics.

To explore the degree of change in TPJ-to-IPS functional connectivity with the subjective degree of intensity of the psychedelic state induced by nitrous oxide, we computed Spearman correlations between TPJ-to-IPS functional connectivity (nitrous oxide versus control condition) and altered states of consciousness score changes (nitrous oxide versus pre-nitrous-oxide baseline). We found that changes in TPJ to right IPS functional connectivity are significantly correlated with five subscales of 11D-altered states questionnaire, including disembodiment (FDR-corrected p=0.018), impaired control and cognition (FDR-corrected p=0.018), anxiety (FDR-corrected p=0.018), changed meaning of percepts (FDR-corrected p=0.019), and experience of unity (FDR-corrected p=0.046) (Figure 6 and Table S6).

**Figure 6:**
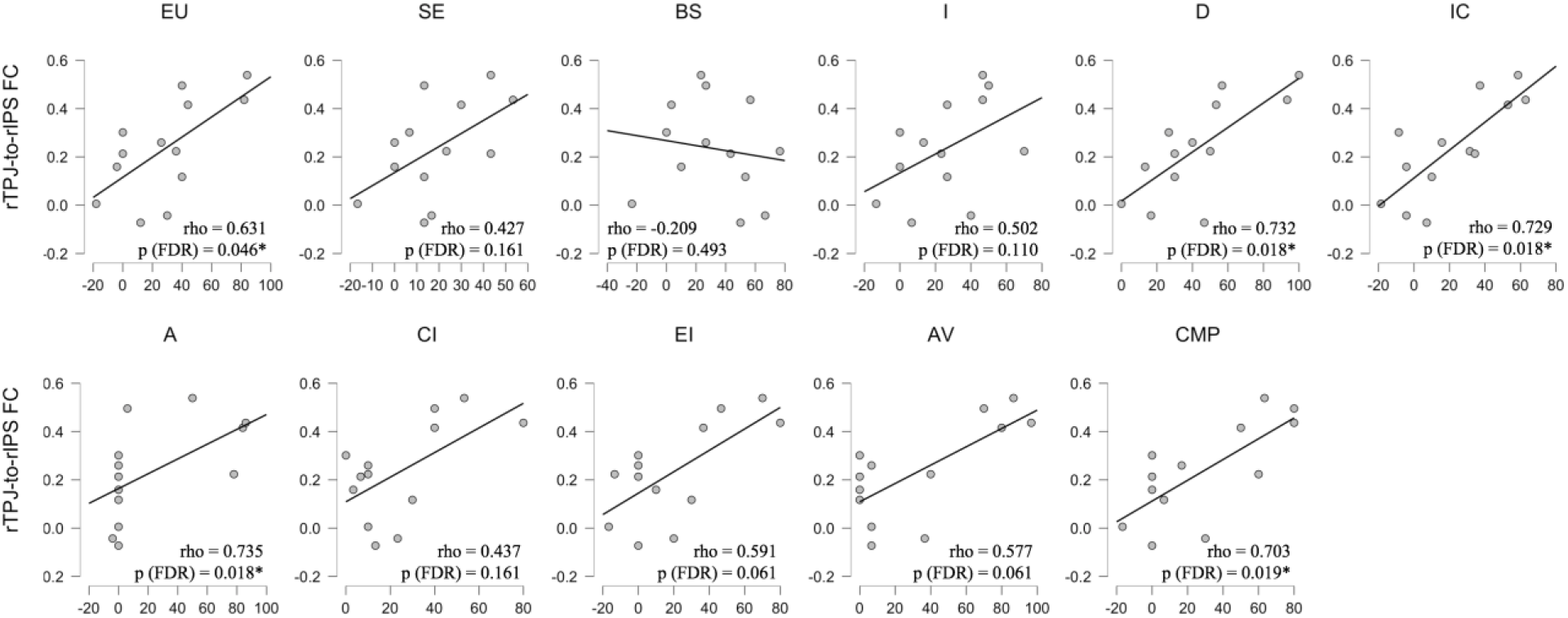
Spearman correlations between right temporoparietal junction to right intraparietal sulcus functional connectivity changes (nitrous oxide versus control condition) and 11D-altered states questionnaire score changes (nitrous oxide versus pre-nitrous oxide baseline). Statistical significance was set at FDR-corrected p < 0.05. EU: experience of unity, SE: spiritual experience, BS: blissful state, I: insightfulness, D: disembodiment, IC: impaired control and cognition, A: anxiety, CI: complex imagery, EI: elementary imagery, AV: audio-visual synesthesia, CMP: changed meaning of percepts.

To complement the functional integration measures of our functional connectivity analysis, we further characterized functional segregation by local correlation analysis. Consistent with our results showing weakened within-network connectivity, we found an overall decrease of local correlation across all three psychedelics; nitrous oxide and ketamine shared some overlap in the right TPJ. In contrast, an overall increase of local correlation was induced by propofol (Figure S2), again distinguishing psychedelic-specific findings from general pharmacological perturbations.

## Discussion

We demonstrate that non-canonical (nitrous oxide, ketamine) and canonical (LSD) psychedelic drugs all reduce within-network functional connectivity and increase between-network connectivity. Common neural correlates induced by these psychedelics, controlled for with the use of a non-psychedelic sedative-hypnotic, included increased connectivity between right TPJ and bilateral IPS and between precuneus and left IPS. These network nodes are located in the posterior hot zone, which has been posited to be critical for content of consciousness (*16*). The consistent results across non-canonical and canonical psychedelics support the hypothesis that there is a common neurobiology underlying the psychedelic effect at the level of large-scale brain networks. Furthermore, the posterior cortical confluence of sensory and association cortex is a biologically plausible candidate for the altered subjective experiences induced by psychedelic drugs.

Specifically, TPJ was the region most consistently involved in psychedelic-induced connectivity changes from both ROI-to-ROI and seed-based functional connectivity analyses. It is known that TPJ is important for multisensory integration and body ownership (*14*), modulation of which might contribute to psychedelic phenomenology (*15*). In support of the specificity of these findings, propofol at sedative concentrations induces functional connectivity changes opposite to those produced by psychedelics, namely, enhanced within-network connectivity and reduced between-network connectivity (*17*). Thus, the effects observed in this study are arguably specific to drugs with psychedelic properties.

These findings inform not only psychedelic neuroscience but emerging psychedelic therapy. Nitrous oxide has been found to have anti-depressant effects in patients with treatment-resistant major depressive disorder (*18*). More recently, it has been shown that a 25% concentration of nitrous oxide is as effective as a 50% concentration for treatment-resistant major depression (*19*).The current study informs the network-level events in the brain that occur during exposure to a comparable concentration of nitrous oxide. Furthermore, ketamine, LSD, and other psychedelics have shown promise as anti-depressants (*20*). Identifying the common neural correlates induced by psychedelic drugs may lead to a more comprehensive mechanistic understanding of therapeutic benefits. Our study informs this neurobiology.

There are numerous limitations to this investigation. First, fMRI datasets were derived from different study protocols and institutions, leading to potential heterogeneity. Second, 3T resolution precludes the ability to make meaningful inferences regarding psychedelic effects on subcortical structures, such as those in the brainstem. Third, nitrous oxide was the only drug formally and prospectively studied for psychedelic phenomenology; volunteers participating in the secondary datasets did not have the same assessment. Thus, we must be circumspect in comparing the psychedelic experience across these drug protocols and restrict interpretation to the neural correlates of psychedelic drug administration. Finally, additional psychedelic drugs such as psilocybin, dimethyltryptamine, and methylenedioxymethamphetamine should be investigated for their effects on connectivity in the posterior cortical hot zone.

Despite these limitations, this study is the first to characterize functional connectivity changes during the administration of psychedelic doses of nitrous oxide and, to our knowledge, the first study to identify cortical network reconfigurations that are common to the administration of both canonical and non-canonical psychedelic drugs. Finally, these network alterations occur consistently in a posterior cortical region argued to be critical for the content of consciousness, presenting a neurobiologically plausible set of network nodes that mediate the psychedelic experience.

## Materials and Methods

### Dataset 1: Nitrous oxide

This study was conducted at the University of Michigan Medical School, where Institutional Review Board (IRB, HUM00096321) approval was obtained. The study team carefully discussed risks and benefits with all participants, after which written informed consent was documented. This analysis was part of a clinical study registered with clinicaltrials.gov (NCT03435055); results from the primary study were posted in July 2021.

A total of 16 participants (ages between 20-34 years old, 8 female) completed two fMRI resting-state scans before and during exposure to subanesthetic concentrations (35%) of nitrous oxide. All participants were classified as American Society of Anesthesiologists physical status I, i.e., completely healthy. Drug abuse and history of psychosis were exclusion criteria, among other health-related conditions (see published registry for details: https://www.clinicaltrials.gov/ct2/show/NCT03435055).

Each volunteer participated in two study visits, an initial consent/pre-scan visit and then a scanning visit within three days. During the pre-scan visit, participants were consented and presented with the details of the study protocol and what they would experience during the scanning session. During the scanning visit, each participant first completed a validated altered states of consciousness questionnaire (*12*). Thereafter, fMRI data were collected during placebo (pure oxygen, 20 minutes) followed by inhaled nitrous oxide at subanesthetic concentrations (35%) over 40 minutes. The initial administration of nitrous oxide was completed outside the scanner and participants were allowed at least a 5-minute equilibrium period prior to the start of the resting state scan. Pressure and visual stimuli, related to a protocol assessing analgesic effects (data not presented here), occurred following the resting state scan to acquire the cleanest data possible. Only resting state fMRI data were analyzed for the purpose of this study. After scanning and 30 minutes of recovery from nitrous oxide administration, the altered states questionnaire was administered again.

To maximize safety, nitrous oxide was delivered using MRI-compatible anesthesia machines, and was first administered outside of the scanner, where airway patency and physiological stability were established prior to imaging. At least two fully trained anesthesiologists directed all anesthetic administration. All participants received ondansetron (4-8 mg IV) with an additional dose of dexamethasone (4 mg IV) if needed to prevent nausea and vomiting. In addition, glycoyrrolate (0.42-0.4 mg IV), labetalol (5-10 mg/kg IV), and midazolam (1-2 mg IV) were available to mitigate any side effects, as needed. Standard intraoperative monitors (electrocardiogram, blood pressure, pulse oximetry, capnography) were used throughout the experiment. Participants wore earplugs and headphones during the fMRI scanning.

Data were acquired at Michigan Medicine, University of Michigan, using a 3T Philips Tesla Philips Achieva (Best, Netherlands). Functional images of the whole brain were acquired by a T2* weighted echo-planar sequence with parameters: 48 slices, TR/TE = 2000/30ms, slice thickness = 3 mm, field of view = 200 × 200mm, flip angle = 90°, scan time = 6 minutes. High-resolution anatomical images were also acquired for resting state fMRI co-registration.

### Dataset 2: Ketamine

This dataset has been previously published based on hypotheses and analyses that are distinct from those of the current study (*21*). The investigation was approved by the IRB of Huashan Hospital, Fudan University; informed consent was obtained from all participants. Twelve right-handed participants were recruited (male/female, 7/5; age, 32 to 66 years). The volunteers were American Society of Anesthesiologists physical status I or II, with no history of brain dysfunction, major organ dysfunction, or use of neuropsychiatric drugs.

Ketamine was infused through an intravenous catheter placed into a vein of the left forearm. fMRI scanning was conducted throughout the whole experiment, ranging from 44 to 62 minutes (means ± SD, 54.6 ± 5.9 minutes). A 10-minute baseline conscious condition was first acquired (except for two participants in which baseline condition was for 6 and 11 minutes). Then, 0.05 mg/ kg per minute of ketamine was infused for 10 minutes (0.5 mg/kg in total), and 0.1 mg/kg per minute was infused for another 10 minutes (1.0 mg/kg in total), except for two participants who only received 0.1 mg/kg per minute infusion for 10 minutes. The ketamine infusion was then discontinued and participants regained responsiveness spontaneously. Two certified anesthesiologists were present throughout the study, with resuscitation equipment always available. Participants wore earplugs and headphones during the fMRI scanning.

A Siemens 3T scanner (Siemens MAGNETOM, Germany) with a standard eight-channel head coil was used. Functional images from the whole brain were acquired by a gradient-echo EPI pulse sequence with parameters: 33 slices, TR/TE = 2000/30 ms, slice thickness = 5 mm, field of view = 210 mm, image matrix = 64 × 64, flip angle = 90°, scan time = 10 minutes. High-resolution anatomical images were also acquired for resting state fMRI co-registration. Only the data derived from the subanesthetic—i.e., psychedelic—dosing was analyzed in the current study.

### Dataset 3: LSD

These data were obtained from an open-access database (doi:10.18112/openneuro.ds003059.v1.0.0); 15 participants were recruited. Drug abuse and history of psychosis were exclusion criteria, among other health-related conditions (see published article for details(*22*)). Volunteers received placebo and LSD across two sessions; the order was counterbalanced across participants. A cannula was inserted and secured in a vein in the antecubital fossa by a medical doctor. All participants received 75 μg of LSD, administered intravenously via a 10ml solution infused over a 2-minute period, followed by an infusion of saline. MRI scanning started approximately 70 minutes after dosing, to capture changes associated with peak intensity between 60 and 90 minutes after administration.

Imaging was performed on a 3T GE HDx system. Functional images across the whole brain were acquired by a gradient-echo EPI pulse sequence with parameters: 35 slices, TR/TE = 2000/35 ms, slice thickness = 3.4 mm, field of view = 220mm, image matrix = 64 × 64, flip angle = 90°, scan time = 7:20 minutes. High-resolution anatomical images were also acquired for resting state fMRI co-registration.

### Dataset 4: Propofol

We used propofol, a sedative-hypnotic drug, as a control for general brain state transitions that are not related to the psychedelic experience. The propofol dataset has been previously published using analyses distinct from those applied here (*21*, *23*, *24*). The study was approved by the IRB of Huashan Hospital, Fudan University. Informed consent was obtained from all participants (n = 26; right-handed; male/female, 12/14; age, 27 to 64 years). Inclusion criteria, anesthetic procedures, fMRI details, scanning parameters, and clinical monitoring were the same as those described for ketamine. Only the data during the subanesthetic dosing (associated with light sedation; n=17) were analyzed in the current study.

Propofol was infused through an intravenous catheter placed in a vein of the right hand or forearm. Propofol was administered using a target-controlled infusion pump to obtain and maintain consistent effect-site concentrations, as estimated by the pharmacokinetic model of propofol (Marsh model). TCI concentrations were increased in 0.1 μg/ml steps beginning at 1.0 μg/ml until reaching the appropriate effect-site concentration. A 5-minute equilibration period was allowed to ensure equilibration of propofol distribution between compartments. The target-controlled propofol infusion was maintained at a stable effect-site concentration for light sedation (1.3 μg/ml). The participants continued to breathe spontaneously with supplemental oxygen via nasal cannula.

A Siemens 3T scanner (Siemens MAGNETOM, Germany) with a standard eight-channel head coil was used. Functional images across the whole brain were acquired by a gradient-echo EPI pulse sequence with parameters: 33 slices, TR/TE = 2000/30 ms, slice thickness = 5 mm, field of view = 210 mm, image matrix = 64 × 64, flip angle = 90°, scan time = 8 minutes. High-resolution anatomical images were also acquired for resting state fMRI co-registration.

### Altered States Questionnaire

The altered-states-of-consciousness questionnaire is composed of 11 dimensions, including the following: experiences of unity, spiritual experience, blissful state, insightfulness, disembodiment, impaired control and cognition, anxiety, complex imagery, elementary imagery, audiovisual synesthesia, and changed meaning of percepts. For all items, the response scale was from 0 (Never) to 10 (Always) with 11 total discrete response options. Scale scores reported here were the average of items within each scale.

### fMRI data preprocessing

Preprocessing steps were implemented in the CONN toolbox (https://web.conn-toolbox.org/) and included: 1) slice timing correction; 2) rigid head motion correction/realignment within and across runs. Frame-wise displacement (FD) of head motion was calculated using frame-wise Euclidean Norm (square root of the sum squares) of the six-dimension motion derivatives. A given frame and its previous frame were tagged as zeros if the frame’s derivative value had a Euclidean Norm above FD = 0.9 mm or the BOLD signal changed above 5 SD (otherwise it was tagged as ones); 3) co-registration with high-resolution anatomical images; 4) spatial normalization into MNI (Montreal Neurological Institute) space and resampling to 3×3×3 mm^3^; 5) time-censored data were low- and high-pass filtered (>0.008, <0.09 Hz). At the same time, various undesired components (e.g., physiological estimates, motion parameters) were removed via linear regression. The undesired components included linear and nonlinear drift, time series of head motion and its temporal derivative, and mean time series from the white matter and cerebrospinal fluid; 6) spatial smoothing with 6 mm full-width at half-maximum isotropic Gaussian kernel. After preprocessing and denoising, the data were visually examined for quality assurance.

Dataset 3 has been preprocessed and published (doi:10.18112/openneuro.ds003059.v1.0.0). The data preprocessing steps included: 1) removal of the first three volumes; 2) de-spiking; 3) slice time correction; 4) motion correction; 5) brain extraction; 6) rigid body registration to anatomical scans; 7) non-linear registration to 2mm MNI brain; 8) scrubbing (Power et al., 2012), using an FD threshold of 0.4 (the mean percentage of volumes scrubbed for placebo and LSD was 0.4 ±0.8% and 1.7 ±2.3%, respectively). The maximum number of scrubbed volumes per scan was 7.1% and scrubbed volumes were replaced with the mean of the surrounding volumes. Additional pre-processing steps included: 9) spatial smoothing of 6mm; 10) band-pass filtering between 0.01 to 0.08 Hz; 11) linear and quadratic de-trending; 12) regressing out undesired components (e.g., motion-related and anatomically related parameters).

### Analysis of ROI-to-ROI Functional Connectivity

Region-of-interest (ROI)-to-ROI functional connectivity analysis was performed using the CONN toolbox (https://web.conn-toolbox.org/). The acquired functional connectivity matrices characterized the connectivity between all pairs of ROIs among a default CONN network parcellation from independent component analysis of the human connectome project (HCP) dataset (n = 497). This HCP-ICA atlas(*25*) covered the main functions of the whole brain, which is divided into seven cerebral networks (default mode network, dorsal attention network, frontoparietal network, language network, salience network, sensorimotor network, visual network) plus one cerebellar network and their corresponding 32 ROIs (Table S1). Each element in the matrix indicates a Fisher-transformed bivariate correlation coefficient between a pair of ROI time courses.

### Analysis of Seed-Based Functional Connectivity

Seed-based functional connectivity maps were computed as the Fisher-transformed bivariate correlation coefficients between the seed BOLD timeseries and each individual voxel BOLD timeseries. Random Field Theory parametric statistics were performed to control for family-wise error at the level of individual clusters.(*26*) The right temporoparietal junction (TPJ) ROI was defined by canonical literature,(*27*) because of its association with psychedelic drug administration in this study, and because it is thought to be critical to multisensory integration, consciousness, and body ownership (*14*, *15*).

### Analysis of Local Correlation

In general, resting-state analytic approaches can be classified as functional integration or functional segregation (*28*, *29*). Functional integration measures (e.g., seed-based connectivity analysis and ROI-to-ROI connectivity analysis) relate to the functional connections between various brain areas (as described above). In contrast, functional segregation measures, e.g., local correlation, focus on the local and differentiated function of specific brain regions. We analyzed local correlation to complement our functional integration analyses and defined it as the average of Pearson correlation coefficients between the time course of each individual voxel and those in a region of neighboring voxels (kernel size was 8 mm; approximately 27 voxels) (*30*).

### Statistical Analysis

For the altered states questionnaire statistical analysis, we performed paired sample t-tests on mean sub-scale scores between each psychedelic condition and baseline condition. After multiple comparison, the statistical significance was set at FDR-corrected p < 0.05. For resting-state fMRI data, standard second-level statistics derived from CONN were used. Due to the differences in scanner parameters between different psychedelic datasets, only within-group statistics were performed, i.e., each psychedelic condition was compared to its own baseline control condition rather than comparisons across different drugs. In the analysis of ROI-to-ROI functional connectivity, we performed functional network connectivity multivariate parametric statistics (*31*) to control family-wise error at the level of individual clusters. We analyzed the set of connections between all pairs of ROIs in relation to the within- and between-network connectivity, then paired sample t-tests were performed to assess the within-group differences between each psychedelic condition and its control condition. Multivariate parametric statistics for functional network connectivity were used and statistical results were thresholded at FDR-corrected p<0.05. In the analysis of seed-based functional connectivity, standard cluster-level inferences based on Gaussian Random Field theory (*32*) were used. We performed paired sample t-tests on TPJ to whole-brain correlation maps for each psychedelic condition and its control condition; the statistical significance was set at FDR-corrected p < 0.05. To assess the degree of change in functional connectivity with the subjective degree of intensity of the psychedelic state induced by nitrous oxide, Spearman correlations were computed between seed-based functional connectivity changes (nitrous oxide versus control condition) and altered-states-of-consciousness score changes (nitrous oxide versus pre-nitrous oxide baseline); statistical significance was set at FDR-corrected p < 0.05. Statistics were computed with IBM SPSS 22 software (IBM SPSS Statistics for Windows, Version 22.0. Armonk, NY, USA: IBM Corp.). For local correlation analysis, we performed paired t-tests on local correlation maps between each psychedelic condition and its control condition; statistical significance was set at FDR-corrected p < 0.05.

## Funding

This work was funded by National Institutes of Health (Bethesda, Maryland, USA) grants R01-GM111293 (to G.A.M., R.E.H.) and T32-GM103730 (to G.A.M., PI, and Z.H., Fellow).

## Author contributions

Conceptualization: RD, ZH, REH, GAM

Methodology: RD, ZH, AGH, REH, GAM

Investigation: TEL, VT, PP, PEV, EJ, AM, REH, GAM

Visualization: RD, ZH

Supervision: GAM

Writing—original draft: RD, ZH, GAM

Writing—review & editing: All authors

## Competing interests

The authors have no conflicts of interest to declare.

## Data and materials availability

All data needed to evaluate the conclusions in this article are present in the main text and the Supplementary Materials. Additional data related to this paper may be requested from the authors. The LSD dataset is available at Openneuro (doi:10.18112/openneuro.ds003059.v1.0.0)

## SUPPLEMENTARY MATERIAL

**Figure S1:**
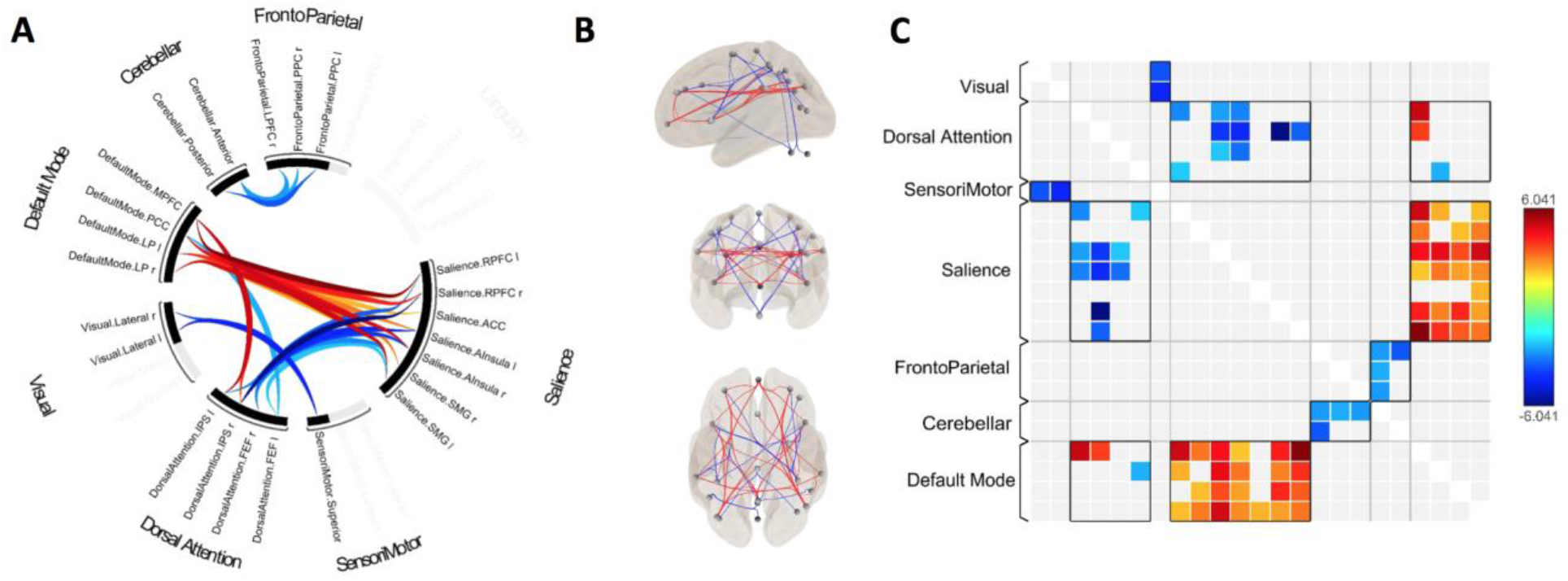
Effects of subanesthetic propofol vs. baseline on functional connectivity. (A) The circle view displays significant functional connectivity changes between ROIs of seven cerebral networks and one cerebellar network. (B) The connectome view displays all of the ROIs with individual suprathreshold connectivity changes lines between them. (C) ROI-to-ROI connectivity changes matrix.

**Figure S2:**
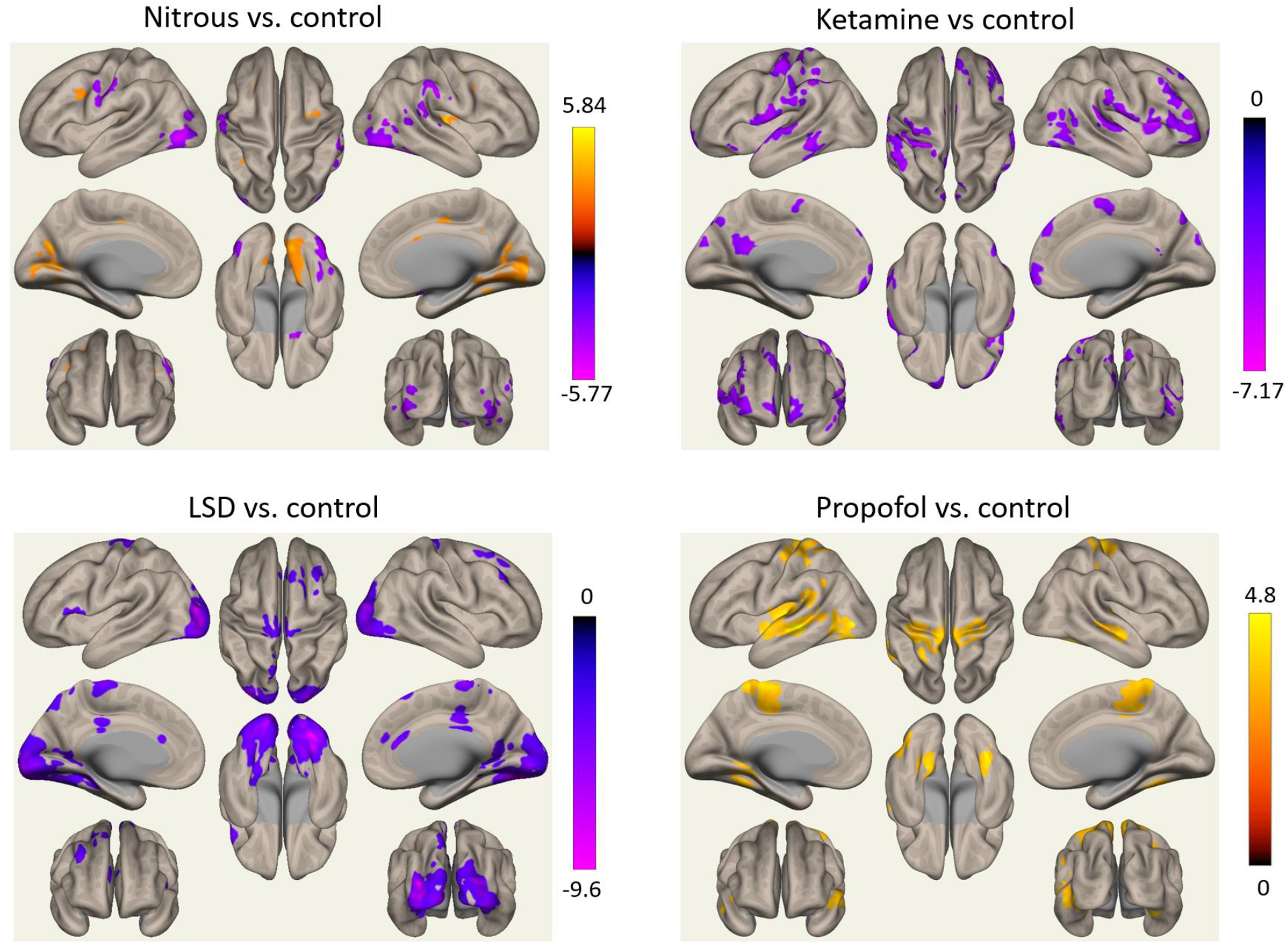
Analysis of local correlation changes between each psychedelic/propofol condition and its control condition.

**Table S1:** Human connectome project/independent component analysis atlas.

| **Network** | **Index** | **Region** | **Hemisphere** | **Coordinate** |
| --- | --- | --- | --- | --- |
| DefaultMode | MPFC | Medial Prefrontal Cortex |  | (1,55,-3) |
| DefaultMode | LP | Lateral Parietal | (L) | (-39,-77,33) |
| DefaultMode | LP | Lateral Parietal | (R) | (47,-67,29) |
| DefaultMode | PCC | Precuneus |  | (1,-61,38) |
| SensoriMotor | Lateral | Lateral SensoriMotor Cortex | (L) | (-55,-12,29) |
| SensoriMotor | Lateral | Lateral SensoriMotor Cortex | (R) | (56,-10,29) |
| SensoriMotor | Superior | Superior SensoriMotor Cortex |  | (0,-31,67) |
| Visual | Medial | Medial Visual Cortex |  | (2,-79,12) |
| Visual | Occipital | Occipital Visual Cortex |  | (0,-93,-4) |
| Visual | Lateral | Lateral Visual Cortex | (L) | (-37,-79,10) |
| Visual | Lateral | Lateral Visual Cortex | (R) | (38,-72,13) |
| Salience | ACC | Anterior Cingulate |  | (0,22,35) |
| Salience | AInsula | Anterior Insula | (L) | (-44,13,1) |
| Salience | AInsula | Anterior Insula | (R) | (47,14,0) |
| Salience | RPFC | Rostral Prefrontal Cortex | (L) | (-32,45,27) |
| Salience | RPFC | Rostral Prefrontal Cortex | (R) | (32,46,27) |
| Salience | SMG | SupraMarginal Gyrus | (L) | (-60,-39,31) |
| Salience | SMG | SupraMarginal Gyrus | (R) | (62,-35,32) |
| DorsalAttention | FEF | Frontal Eye Field | (L) | (-27,-9,64) |
| DorsalAttention | FEF | Frontal Eye Field | (R) | (30,-6,64) |
| DorsalAttention | IPS | IntraParietal Sulcus | (L) | (-39,-43,52) |
| DorsalAttention | IPS | IntraParietal Sulcus | (R) | (39,-42,54) |
| FrontoParietal | LPFC | Lateral Prefrontal Cortex | (L) | (-43,33,28) |
| FrontoParietal | PPC | Posterior Parietal Cortex | (L) | (-46,-58,49) |
| FrontoParietal | LPFC | Lateral Prefrontal Cortex | (R) | (41,38,30) |
| FrontoParietal | PPC | Posterior Parietal Cortex | (R) | (52,-52,45) |
| Language | IFG | Inferior Frontal Gyrus | (L) | (-51,26,2) |
| Language | IFG | Inferior Frontal Gyrus | (R) | (54,28,1) |
| Language | pSTG | Posterior Superior Temporal Gyrus | (L) | (-57,-47,15) |
| Language | pSTG | Posterior Superior Temporal Gyrus | (R) | (59,-42,13) |
| Cerebellar | Anterior | Anterior Cerebellar |  | (0,-63,-30) |
| Cerebellar | Posterior | Posterior Cerebellar |  | (0,-79,-32) |

**Table S2:** Nitrous oxide vs. baseline ROI-to-ROI functional connectivity results.

| Analysis Unit |  | From | Hem | Coordinate | To | Hem | Coordinate | Statistic | p-unc | p-FDR |
| --- | --- | --- | --- | --- | --- | --- | --- | --- | --- | --- |
| Cluster | 1/36 |  |  |  |  |  |  | F(1,15) = 24.1 | 0.000189 | 0.0068 |
|  | Connection | networks.Visual.Lateral | (L) | (-37,-79,10) | -networks.Salience.RPFC | (R) | (32,46,27) | T(15) = 4.81 | 0.00023 | 0.004988 |
|  | Connection | networks.Visual.Lateral | (L) | (-37,-79,10) | -networks.Salience.ACC |  | (0,22,35) | T(15) = 4.64 | 0.000322 | 0.004988 |
|  | Connection | networks.Visual.Lateral | (L) | (-37,-79,10) | -networks.Salience.Alnsula | (R) | (47,14,0) | T(15) = 4.13 | 0.000883 | 0.009123 |
|  | Connection | networks.Visual.Lateral | (R) | (38,-72,13) | -networks.Salience.RPFC | (R) | (32,46,27) | T(15) = 3.97 | 0.001234 | 0.015283 |
|  | Connection | networks.Visual.Lateral | (R) | (38,-72,13) | -networks.Salience.SMG | (R) | (62,-35,32) | T(15) = 3.88 | 0.001479 | 0.015283 |
|  | Connection | networks.Visual.Lateral | (R) | (38,-72,13) | -networks.Salience.Alnsula | (R) | (47,14,0) | T(15) = 3.31 | 0.004739 | 0.021294 |
|  | Connection | networks.Visual.Lateral | (R) | (38,-72,13) | -networks.Salience.SMG | (L) | (-60,-39,31) | T(15) = 3.3 | 0.004812 | 0.021294 |
|  | Connection | networks.Visual.Lateral | (R) | (38,-72,13) | -networks.Salience.ACC |  | (0,22,35) | T(15) = 3.2 | 0.005913 | 0.021294 |
|  | Connection | networks.Visual.Lateral | (L) | (-37,-79,10) | -networks.Salience.RPFC | (L) | (-32,45,27) | T(15) = 2.79 | 0.013756 | 0.06092 |
|  | Connection | networks.Visual.Lateral | (R) | (38,-72,13) | -networks.Salience.RPFC | (L) | (-32,45,27) | T(15) = 2.4 | 0.029775 | 0.076919 |
|  | Connection | networks.Visual.Lateral | (L) | (-37,-79,10) | -networks.Salience.Alnsula | (L) | (-44,13,1) | T(15) = 2.51 | 0.023805 | 0.08781 |
|  | Connection | networks.Visual.Lateral | (L) | (-37,-79,10) | -networks.Salience.SMG | (R) | (62,-35,32) | T(15) = 2.47 | 0.026252 | 0.08781 |
|  | Connection | networks.Visual.Lateral | (L) | (-37,-79,10) | -networks.Salience.SMG | (L) | (-60,-39,31) | T(15) = 2.43 | 0.028326 | 0.08781 |
|  | Connection | networks.Visual.Occipital |  | (0,-93,-4) | -networks.Salience.RPFC | (L) | (-32,45,27) | T(15) = 2.69 | 0.016813 | 0.291383 |
|  | Connection | networks.Visual.Occipital |  | (0,-93,-4) | -networks.Salience.RPFC | (R) | (32,46,27) | T(15) = 2.44 | 0.027764 | 0.291383 |
| Cluster | 2/36 |  |  |  |  |  |  | F(1,15) = 13.56 | 0.002219 | 0.024456 |
|  | Connection | networks.DorsalAttention.FEF | (L) | (-27,-9,64) | -networks.FrontoParietal.LPFC | (R) | (41,38,30) | T(15) = 3.66 | 0.002317 | 0.07182 |
|  | Connection | networks.DorsalAttention.IPS | (L) | (-39,-43,52) | -networks.FrontoParietal.LPFC | (R) | (41,38,30) | T(15) = 3.28 | 0.005104 | 0.158227 |
|  | Connection | networks.DorsalAttention.IPS | (L) | (-39,-43,52) | -networks.FrontoParietal.PPC | (R) | (52,-52,45) | T(15) = 2.91 | 0.010689 | 0.165675 |
|  | Connection | networks.DorsalAttention.IPS | (R) | (39,-42,54) | -networks.FrontoParietal.PPC | (R) | (52,-52,45) | T(15) = 2.48 | 0.025576 | 0.181394 |
|  | Connection | networks.DorsalAttention.FEF | (L) | (-27,-9,64) | -networks.FrontoParietal.LPFC | (L) | (-43,33,28) | T(15) = 2.34 | 0.03379 | 0.448816 |
| Cluster | 3/36 |  |  |  |  |  |  | F(1,15) = 12.54 | 0.002958 | 0.024456 |
|  | Connection | networks.Salience.Alnsula | (R) | (47,14,0) | -networks.Salience.RPFC | (R) | (32,46,27) | T(15) = -2.97 | 0.009542 | 0.073951 |
|  | Connection | networks.Salience.RPFC | (R) | (32,46,27) | -networks.Salience.RPFC | (L) | (-32,45,27) | T(15) = -2.54 | 0.022541 | 0.099824 |
|  | Connection | networks.Salience.Alnsula | (L) | (-44,13,1) | -networks.Salience.RPFC | (L) | (-32,45,27) | T(15) = -3.08 | 0.007697 | 0.11931 |
|  | Connection | networks.Salience.Alnsula | (L) | (-44,13,1) | -networks.Salience.RPFC | (R) | (32,46,27) | T(15) = -2.59 | 0.020533 | 0.122994 |
|  | Connection | networks.Salience.ACC |  | (0,22,35) | -networks.Salience.RPFC | (R) | (32,46,27) | T(15) = -2.17 | 0.046201 | 0.143728 |
|  | Connection | networks.Salience.Alnsula | (R) | (47,14,0) | -networks.Salience.RPFC | (L) | (-32,45,27) | T(15) = -2.16 | 0.047029 | 0.218477 |
| Cluster | 4/36 |  |  |  |  |  |  | F(1,15) = 12 | 0.003466 | 0.024456 |
|  | Connection | networks.SensoriMotor.Lateral | (L) | (-55,-12,29) | -networks.Language.IFG | (L) | (-51,26,2) | T(15) = 4.1 | 0.00095 | 0.029438 |
|  | Connection | networks.SensoriMotor.Lateral | (L) | (-55,-12,29) | -networks.Language.pSTG | (L) | (-57,-47,15) | T(15) = 2.96 | 0.009799 | 0.060755 |
|  | Connection | networks.SensoriMotor.Lateral | (L) | (-55,-12,29) | -networks.Language.pSTG | (R) | (59,-42,13) | T(15) = 2.5 | 0.024652 | 0.095526 |
|  | Connection | networks.SensoriMotor.Superior |  | (0,-31,67) | -networks.Language.IFG | (R) | (54,28,1) | T(15) = 2.73 | 0.015372 | 0.116259 |
|  | Connection | networks.SensoriMotor.Lateral | (R) | (56,-10,29) | -networks.Language.IFG | (L) | (-51,26,2) | T(15) = 2.84 | 0.012332 | 0.127428 |
|  | Connection | networks.SensoriMotor.Lateral | (R) | (56,-10,29) | -networks.Language.pSTG | (R) | (59,-42,13) | T(15) = 2.62 | 0.019401 | 0.150361 |
|  | Connection | networks.SensoriMotor.Lateral | (R) | (56,-10,29) | -networks.Language.pSTG | (L) | (-57,-47,15) | T(15) = 2.25 | 0.039604 | 0.245548 |
| Cluster | 5/36 |  |  |  |  |  |  | F(1,15) = 11.01 | 0.004675 | 0.024456 |
|  | Connection | networks.DorsalAttention.IPS | (R) | (39,-42,54) | -networks.Language.pSTG | (R) | (59,-42,13) | T(15) = 3.71 | 0.00211 | 0.032711 |
|  | Connection | networks.DorsalAttention.IPS | (L) | (-39,-43,52) | -networks.Language.pSTG | (L) | (-57,-47,15) | T(15) = 2.43 | 0.02811 | 0.174282 |
|  | Connection | networks.DorsalAttention.IPS | (R) | (39,-42,54) | -networks.Language.pSTG | (L) | (-57,-47,15) | T(15) = 2.38 | 0.031055 | 0.181394 |
| Cluster | 6/36 |  |  |  |  |  |  | F(1,15) = 10.83 | 0.004949 | 0.024456 |
|  | Connection | networks.Language.IFG | (R) | (54,28,1) | -networks.Language.pSTG | (L) | (-57,-47,15) | T(15) = -3.14 | 0.006716 | 0.106679 |
|  | Connection | networks.Language.IFG | (R) | (54,28,1) | -networks.Language.pSTG | (R) | (59,-42,13) | T(15) = -2.62 | 0.019478 | 0.150954 |
|  | Connection | networks.Language.IFG | (R) | (54,28,1) | -networks.Language.IFG | (L) | (-51,26,2) | T(15) = -2.45 | 0.027283 | 0.169152 |
| Cluster | 7/36 |  |  |  |  |  |  | F(1,15) = 10.61 | 0.005302 | 0.024456 |
|  | Connection | networks.Salience.RPFC | (L) | (-32,45,27) | -networks.DefaultMode.LP | (L) | (-39,-77,33) | T(15) = 4.59 | 0.000351 | 0.010888 |
|  | Connection | networks.Salience.Alnsula | (L) | (-44,13,1) | -networks.DefaultMode.LP | (R) | (47,-67,29) | T(15) = 3.69 | 0.002162 | 0.067008 |
|  | Connection | networks.Salience.ACC |  | (0,22,35) | -networks.DefaultMode.LP | (R) | (47,-67,29) | T(15) = 2.6 | 0.020165 | 0.089302 |
|  | Connection | networks.Salience.RPFC | (L) | (-32,45,27) | -networks.DefaultMode.PCC |  | (1,-61,38) | T(15) = 2.61 | 0.01987 | 0.116461 |
|  | Connection | networks.Salience.SMG | (L) | (-60,-39,31) | -networks.DefaultMode.PCC |  | (1,-61,38) | T(15) = 2.88 | 0.011336 | 0.117143 |
|  | Connection | networks.Salience.SMG | (L) | (-60,-39,31) | -networks.DefaultMode.MPFC |  | (1,55,-3) | T(15) = 2.74 | 0.015334 | 0.118836 |
|  | Connection | networks.Salience.Alnsula | (L) | (-44,13,1) | -networks.DefaultMode.PCC |  | (1,-61,38) | T(15) = 2.15 | 0.04807 | 0.187998 |
| Cluster | 8/36 |  |  |  |  |  |  | F(1,15) = 10.53 | 0.005435 | 0.024456 |
|  | Connection | networks.DorsalAttention.IPS | (R) | (39,-42,54) | -networks.DefaultMode.LP | (R) | (47,-67,29) | T(15) = 4.05 | 0.001045 | 0.032385 |
|  | Connection | networks.DorsalAttention.IPS | (L) | (-39,-43,52) | -networks.DefaultMode.PCC |  | (1,-61,38) | T(15) = 2.62 | 0.01933 | 0.174282 |
|  | Connection | networks.DorsalAttention.IPS | (L) | (-39,-43,52) | -networks.DefaultMode.LP | (R) | (47,-67,29) | T(15) = 2.45 | 0.026974 | 0.174282 |
|  | Connection | networks.DorsalAttention.IPS | (R) | (39,-42,54) | -networks.DefaultMode.LP | (L) | (-39,-77,33) | T(15) = 2.32 | 0.035108 | 0.181394 |
|  | Connection | networks.DorsalAttention.IPS | (R) | (39,-42,54) | -networks.DefaultMode.PCC |  | (1,-61,38) | T(15) = 2.2 | 0.043538 | 0.192809 |

**Table S3:**
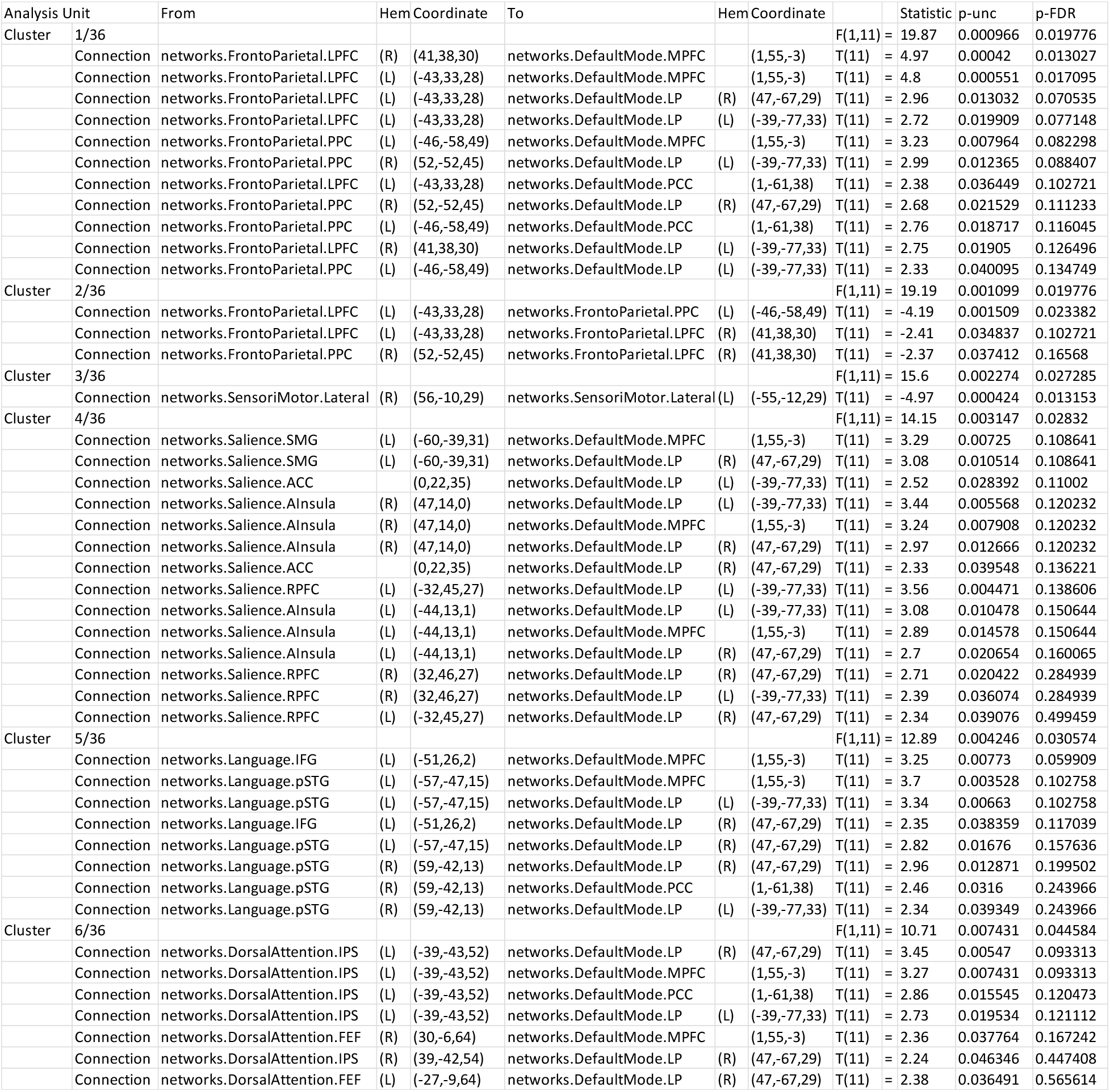
Ketamine vs. baseline ROI-to-ROI functional connectivity results.

**Table S4:**
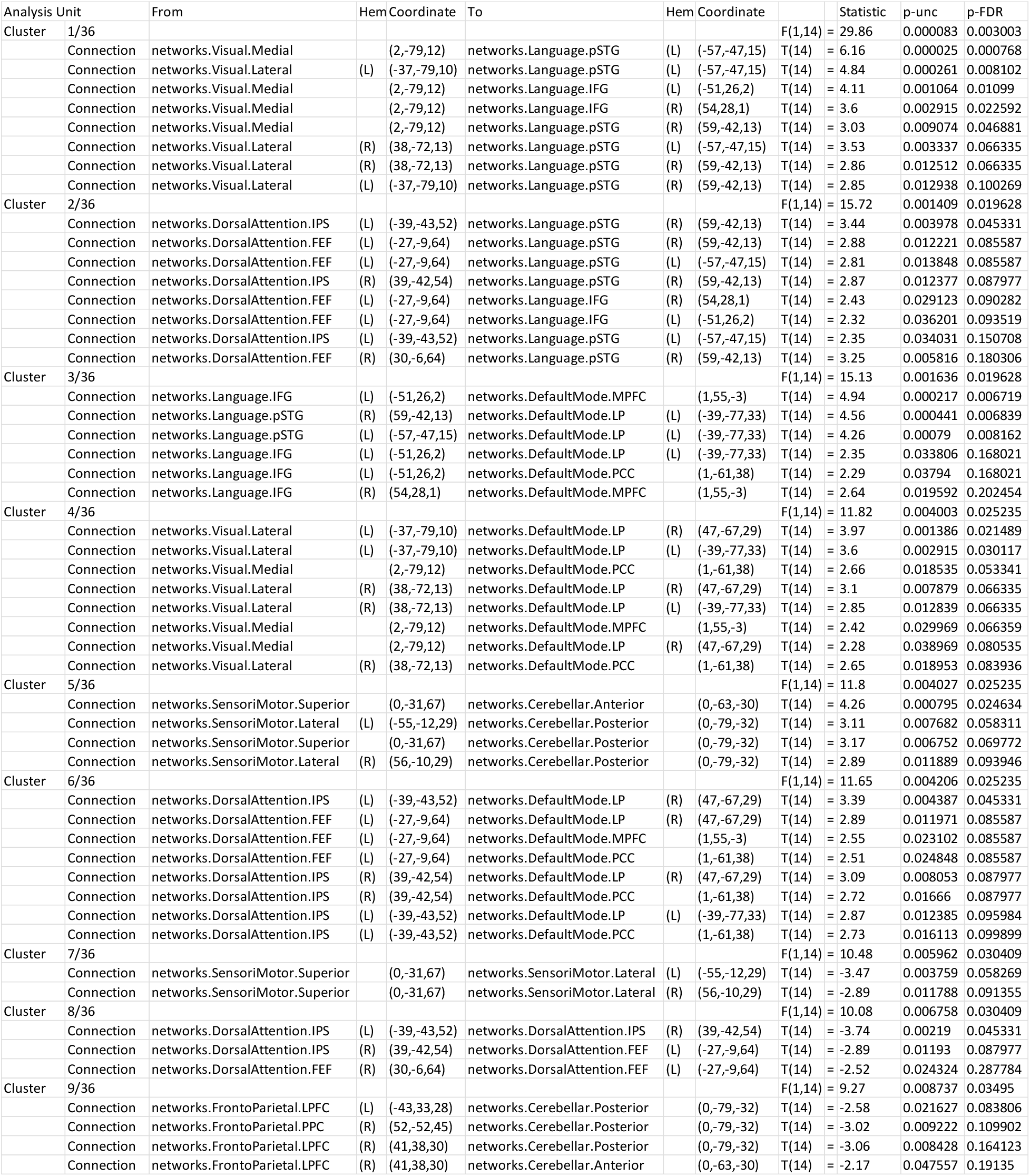
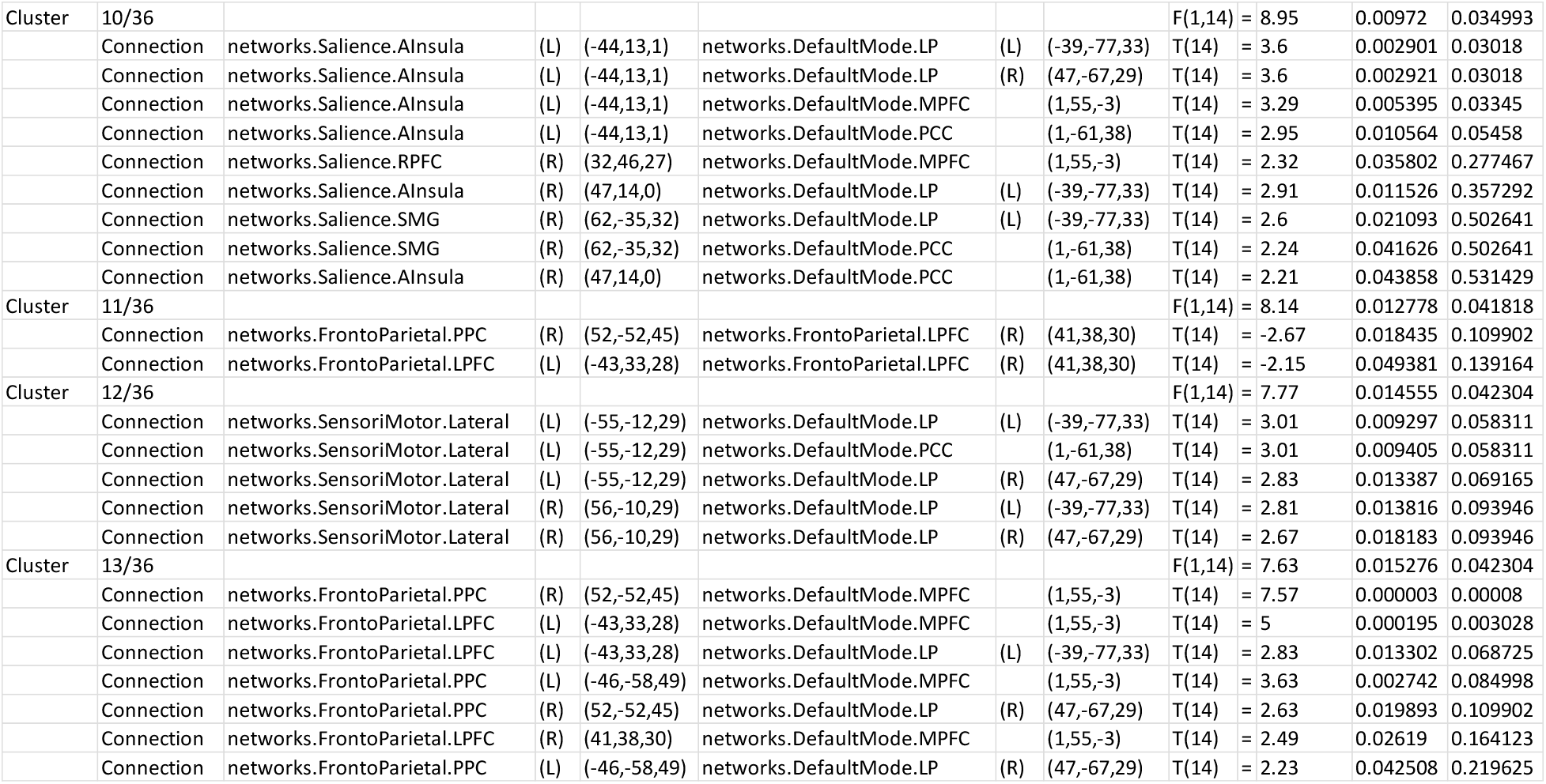
LSD vs. placebo ROI-to-ROI functional connectivity results.

| Analysis Unit | From | Hem Coordinate | To | Hem Coordinate | Statistic | p-unc | p-FDR |
| --- | --- | --- | --- | --- | --- | --- | --- |
| Cluster 1/36 |  |  |  |  | F(1,14) = 29.86 | 0.000083 | 0.003003 |
| Connection | networks.Visual.Medial | (2,-79,12) | networks.Language.pSTG | (L) (-57,-47,15) | T(14) = 6.16 | 0.000025 | 0.000768 |
| Connection | networks.Visual.Lateral | (L) (-37,-79,10) | networks.Language.pSTG | (L) (-57,-47,15) | T(14) = 4.84 | 0.000261 | 0.008102 |
| Connection | networks.Visual.Medial | (2,-79,12) | networks.Language.IFG | (L) (-51,26,2) | T(14) = 4.11 | 0.001064 | 0.01099 |
| Connection | networks.Visual.Medial | (2,-79,12) | networks.Language.IFG | (R) (54,28,1) | T(14) = 3.6 | 0.002915 | 0.022592 |
| Connection | networks.Visual.Medial | (2,-79,12) | networks.Language.pSTG | (R) (59,-42,13) | T(14) = 3.03 | 0.009074 | 0.046881 |
| Connection | networks.Visual.Lateral | (R) (38,-72,13) | networks.Language.pSTG | (L) (-57,-47,15) | T(14) = 3.53 | 0.003337 | 0.066335 |
| Connection | networks.Visual.Lateral | (R) (38,-72,13) | networks.Language.pSTG | (R) (59,-42,13) | T(14) = 2.86 | 0.012512 | 0.066335 |
| Connection | networks.Visual.Lateral | (L) (-37,-79,10) | networks.Language.pSTG | (R) (59,-42,13) | T(14) = 2.85 | 0.012938 | 0.100269 |
| Cluster 2/36 |  |  |  |  | F(1,14) = 15.72 | 0.001409 | 0.019628 |
| Connection | networks.DorsalAttention.IPS | (L) (-39,-43,52) | networks.Language.pSTG | (R) (59,-42,13) | T(14) = 3.44 | 0.003978 | 0.045331 |
| Connection | networks.DorsalAttention.FEF | (L) (-27,-9,64) | networks.Language.pSTG | (R) (59,-42,13) | T(14) = 2.88 | 0.012221 | 0.085587 |
| Connection | networks.DorsalAttention.FEF | (L) (-27,-9,64) | networks.Language.pSTG | (L) (-57,-47,15) | T(14) = 2.81 | 0.013848 | 0.085587 |
| Connection | networks.DorsalAttention.IPS | (R) (39,-42,54) | networks.Language.pSTG | (R) (59,-42,13) | T(14) = 2.87 | 0.012377 | 0.087977 |
| Connection | networks.DorsalAttention.FEF | (L) (-27,-9,64) | networks.Language.IFG | (R) (54,28,1) | T(14) = 2.43 | 0.029123 | 0.090282 |
| Connection | networks.DorsalAttention.FEF | (L) (-27,-9,64) | networks.Language.IFG | (L) (-51,26,2) | T(14) = 2.32 | 0.036201 | 0.093519 |
| Connection | networks.DorsalAttention.IPS | (L) (-39,-43,52) | networks.Language.pSTG | (L) (-57,-47,15) | T(14) = 2.35 | 0.034031 | 0.150708 |
| Connection | networks.DorsalAttention.FEF | (R) (30,-6,64) | networks.Language.pSTG | (R) (59,-42,13) | T(14) = 3.25 | 0.005816 | 0.180306 |
| Cluster 3/36 |  |  |  |  | F(1,14) = 15.13 | 0.001636 | 0.019628 |
| Connection | networks.Language.IFG | (L) (-51,26,2) | networks.DefaultMode.MPFC | (1,55,-3) | T(14) = 4.94 | 0.000217 | 0.006719 |
| Connection | networks.Language.pSTG | (R) (59,-42,13) | networks.DefaultMode.LP | (L) (-39,-77,33) | T(14) = 4.56 | 0.000441 | 0.006839 |
| Connection | networks.Language.pSTG | (L) (-57,-47,15) | networks.DefaultMode.LP | (L) (-39,-77,33) | T(14) = 4.26 | 0.00079 | 0.008162 |
| Connection | networks.Language.IFG | (L) (-51,26,2) | networks.DefaultMode.LP | (L) (-39,-77,33) | T(14) = 2.35 | 0.033806 | 0.168021 |
| Connection | networks.Language.IFG | (L) (-51,26,2) | networks.DefaultMode.PCC | (1,-61,38) | T(14) = 2.29 | 0.03794 | 0.168021 |
| Connection | networks.Language.IFG | (R) (54,28,1) | networks.DefaultMode.MPFC | (1,55,-3) | T(14) = 2.64 | 0.019592 | 0.202454 |
| Cluster 4/36 |  |  |  |  | F(1,14) = 11.82 | 0.004003 | 0.025235 |
| Connection | networks.Visual.Lateral | (L) (-37,-79,10) | networks.DefaultMode.LP | (R) (47,-67,29) | T(14) = 3.97 | 0.001386 | 0.021489 |
| Connection | networks.Visual.Lateral | (L) (-37,-79,10) | networks.DefaultMode.LP | (L) (-39,-77,33) | T(14) = 3.6 | 0.002915 | 0.030117 |
| Connection | networks.Visual.Medial | (2,-79,12) | networks.DefaultMode.PCC | (1,-61,38) | T(14) = 2.66 | 0.018535 | 0.053341 |
| Connection | networks.Visual.Lateral | (R) (38,-72,13) | networks.DefaultMode.LP | (R) (47,-67,29) | T(14) = 3.1 | 0.007879 | 0.066335 |
| Connection | networks.Visual.Lateral | (R) (38,-72,13) | networks.DefaultMode.LP | (L) (-39,-77,33) | T(14) = 2.85 | 0.012839 | 0.066335 |
| Connection | networks.Visual.Medial | (2,-79,12) | networks.DefaultMode.MPFC | (1,55,-3) | T(14) = 2.42 | 0.029969 | 0.066359 |
| Connection | networks.Visual.Medial | (2,-79,12) | networks.DefaultMode.LP | (R) (47,-67,29) | T(14) = 2.28 | 0.038969 | 0.080535 |
| Connection | networks.Visual.Lateral | (R) (38,-72,13) | networks.DefaultMode.PCC | (1,-61,38) | T(14) = 2.65 | 0.018953 | 0.083936 |
| Cluster 5/36 |  |  |  |  | F(1,14) = 11.8 | 0.004027 | 0.025235 |
| Connection | networks.SensoriMotor.Superior | (0,-31,67) | networks.Cerebellar.Anterior | (0,-63,-30) | T(14) = 4.26 | 0.000795 | 0.024634 |
| Connection | networks.SensoriMotor.Lateral | (L) (-55,-12,29) | networks.Cerebellar.Posterior | (0,-79,-32) | T(14) = 3.11 | 0.007682 | 0.058311 |
| Connection | networks.SensoriMotor.Superior | (0,-31,67) | networks.Cerebellar.Posterior | (0,-79,-32) | T(14) = 3.17 | 0.006752 | 0.069772 |
| Connection | networks.SensoriMotor.Lateral | (R) (56,-10,29) | networks.Cerebellar.Posterior | (0,-79,-32) | T(14) = 2.89 | 0.011889 | 0.093946 |
| Cluster 6/36 |  |  |  |  | F(1,14) = 11.65 | 0.004206 | 0.025235 |
| Connection | networks.DorsalAttention.IPS | (L) (-39,-43,52) | networks.DefaultMode.LP | (R) (47,-67,29) | T(14) = 3.39 | 0.004387 | 0.045331 |
| Connection | networks.DorsalAttention.FEF | (L) (-27,-9,64) | networks.DefaultMode.LP | (R) (47,-67,29) | T(14) = 2.89 | 0.011971 | 0.085587 |
| Connection | networks.DorsalAttention.FEF | (L) (-27,-9,64) | networks.DefaultMode.MPFC | (1,55,-3) | T(14) = 2.55 | 0.023102 | 0.085587 |
| Connection | networks.DorsalAttention.FEF | (L) (-27,-9,64) | networks.DefaultMode.PCC | (1,-61,38) | T(14) = 2.51 | 0.024848 | 0.085587 |
| Connection | networks.DorsalAttention.IPS | (R) (39,-42,54) | networks.DefaultMode.LP | (R) (47,-67,29) | T(14) = 3.09 | 0.008053 | 0.087977 |
| Connection | networks.DorsalAttention.IPS | (R) (39,-42,54) | networks.DefaultMode.PCC | (1,-61,38) | T(14) = 2.72 | 0.01666 | 0.087977 |
| Connection | networks.DorsalAttention.IPS | (L) (-39,-43,52) | networks.DefaultMode.LP | (L) (-39,-77,33) | T(14) = 2.87 | 0.012385 | 0.095984 |
| Connection | networks.DorsalAttention.IPS | (L) (-39,-43,52) | networks.DefaultMode.PCC | (1,-61,38) | T(14) = 2.73 | 0.016113 | 0.099899 |
| Cluster 7/36 |  |  |  |  | F(1,14) = 10.48 | 0.005962 | 0.030409 |
| Connection | networks.SensoriMotor.Superior | (0,-31,67) | networks.SensoriMotor.Lateral | (L) (-55,-12,29) | T(14) = -3.47 | 0.003759 | 0.058269 |
| Connection | networks.SensoriMotor.Superior | (0,-31,67) | networks.SensoriMotor.Lateral | (R) (56,-10,29) | T(14) = -2.89 | 0.011788 | 0.091355 |
| Cluster 8/36 |  |  |  |  | F(1,14) = 10.08 | 0.006758 | 0.030409 |
| Connection | networks.DorsalAttention.IPS | (L) (-39,-43,52) | networks.DorsalAttention.IPS | (R) (39,-42,54) | T(14) = -3.74 | 0.00219 | 0.045331 |
| Connection | networks.DorsalAttention.IPS | (R) (39,-42,54) | networks.DorsalAttention.FEF | (L) (-27,-9,64) | T(14) = -2.89 | 0.01193 | 0.087977 |
| Connection | networks.DorsalAttention.FEF | (R) (30,-6,64) | networks.DorsalAttention.FEF | (L) (-27,-9,64) | T(14) = -2.52 | 0.024324 | 0.287784 |
| Cluster 9/36 |  |  |  |  | F(1,14) = 9.27 | 0.008737 | 0.03495 |
| Connection | networks.FrontoParietal.LPFC | (L) (-43,33,28) | networks.Cerebellar.Posterior | (0,-79,-32) | T(14) = -2.58 | 0.021627 | 0.083806 |
| Connection | networks.FrontoParietal.PPC | (R) (52,-52,45) | networks.Cerebellar.Posterior | (0,-79,-32) | T(14) = -3.02 | 0.009222 | 0.109902 |
| Connection | networks.FrontoParietal.LPFC | (R) (41,38,30) | networks.Cerebellar.Posterior | (0,-79,-32) | T(14) = -3.06 | 0.008428 | 0.164123 |
| Connection | networks.FrontoParietal.LPFC | (R) (41,38,30) | networks.Cerebellar.Anterior | (0,-63,-30) | T(14) = -2.17 | 0.047557 | 0.19135 |

|  |  |  |  |  |  |  |  |  |  |  |
| --- | --- | --- | --- | --- | --- | --- | --- | --- | --- | --- |
| Cluster | 10/36 |  |  |  |  |  |  | F(1,14) = 8.95 | 0.00972 | 0.034993 |
|  | Connection | networks.Salience.AInsula | (L) | (-44,13,1) | networks.DefaultMode.LP | (L) | (-39,-77,33) | T(14) = 3.6 | 0.002901 | 0.03018 |
|  | Connection | networks.Salience.AInsula | (L) | (-44,13,1) | networks.DefaultMode.LP | (R) | (47,-67,29) | T(14) = 3.6 | 0.002921 | 0.03018 |
|  | Connection | networks.Salience.AInsula | (L) | (-44,13,1) | networks.DefaultMode.MPFC |  | (1,55,-3) | T(14) = 3.29 | 0.005395 | 0.03345 |
|  | Connection | networks.Salience.AInsula | (L) | (-44,13,1) | networks.DefaultMode.PCC |  | (1,-61,38) | T(14) = 2.95 | 0.010564 | 0.05458 |
|  | Connection | networks.Salience.RPFC | (R) | (32,46,27) | networks.DefaultMode.MPFC |  | (1,55,-3) | T(14) = 2.32 | 0.035802 | 0.277467 |
|  | Connection | networks.Salience.AInsula | (R) | (47,14,0) | networks.DefaultMode.LP | (L) | (-39,-77,33) | T(14) = 2.91 | 0.011526 | 0.357292 |
|  | Connection | networks.Salience.SMG | (R) | (62,-35,32) | networks.DefaultMode.LP | (L) | (-39,-77,33) | T(14) = 2.6 | 0.021093 | 0.502641 |
|  | Connection | networks.Salience.SMG | (R) | (62,-35,32) | networks.DefaultMode.PCC |  | (1,-61,38) | T(14) = 2.24 | 0.041626 | 0.502641 |
|  | Connection | networks.Salience.AInsula | (R) | (47,14,0) | networks.DefaultMode.PCC |  | (1,-61,38) | T(14) = 2.21 | 0.043858 | 0.531429 |
| Cluster | 11/36 |  |  |  |  |  |  | F(1,14) = 8.14 | 0.012778 | 0.041818 |
|  | Connection | networks.FrontoParietal.PPC | (R) | (52,-52,45) | networks.FrontoParietal.LPFC | (R) | (41,38,30) | T(14) = -2.67 | 0.018435 | 0.109902 |
|  | Connection | networks.FrontoParietal.LPFC | (L) | (-43,33,28) | networks.FrontoParietal.LPFC | (R) | (41,38,30) | T(14) = -2.15 | 0.049381 | 0.139164 |
| Cluster | 12/36 |  |  |  |  |  |  | F(1,14) = 7.77 | 0.014555 | 0.042304 |
|  | Connection | networks.SensoriMotor.Lateral | (L) | (-55,-12,29) | networks.DefaultMode.LP | (L) | (-39,-77,33) | T(14) = 3.01 | 0.009297 | 0.058311 |
|  | Connection | networks.SensoriMotor.Lateral | (L) | (-55,-12,29) | networks.DefaultMode.PCC |  | (1,-61,38) | T(14) = 3.01 | 0.009405 | 0.058311 |
|  | Connection | networks.SensoriMotor.Lateral | (L) | (-55,-12,29) | networks.DefaultMode.LP | (R) | (47,-67,29) | T(14) = 2.83 | 0.013387 | 0.069165 |
|  | Connection | networks.SensoriMotor.Lateral | (R) | (56,-10,29) | networks.DefaultMode.LP | (L) | (-39,-77,33) | T(14) = 2.81 | 0.013816 | 0.093946 |
|  | Connection | networks.SensoriMotor.Lateral | (R) | (56,-10,29) | networks.DefaultMode.LP | (R) | (47,-67,29) | T(14) = 2.67 | 0.018183 | 0.093946 |
| Cluster | 13/36 |  |  |  |  |  |  | F(1,14) = 7.63 | 0.015276 | 0.042304 |
|  | Connection | networks.FrontoParietal.PPC | (R) | (52,-52,45) | networks.DefaultMode.MPFC |  | (1,55,-3) | T(14) = 7.57 | 0.000003 | 0.00008 |
|  | Connection | networks.FrontoParietal.LPFC | (L) | (-43,33,28) | networks.DefaultMode.MPFC |  | (1,55,-3) | T(14) = 5 | 0.000195 | 0.003028 |
|  | Connection | networks.FrontoParietal.LPFC | (L) | (-43,33,28) | networks.DefaultMode.LP | (L) | (-39,-77,33) | T(14) = 2.83 | 0.013302 | 0.068725 |
|  | Connection | networks.FrontoParietal.PPC | (L) | (-46,-58,49) | networks.DefaultMode.MPFC |  | (1,55,-3) | T(14) = 3.63 | 0.002742 | 0.084998 |
|  | Connection | networks.FrontoParietal.PPC | (R) | (52,-52,45) | networks.DefaultMode.LP | (R) | (47,-67,29) | T(14) = 2.63 | 0.019893 | 0.109902 |
|  | Connection | networks.FrontoParietal.LPFC | (R) | (41,38,30) | networks.DefaultMode.MPFC |  | (1,55,-3) | T(14) = 2.49 | 0.02619 | 0.164123 |
|  | Connection | networks.FrontoParietal.PPC | (L) | (-46,-58,49) | networks.DefaultMode.LP | (R) | (47,-67,29) | T(14) = 2.23 | 0.042508 | 0.219625 |

**Table S5:**
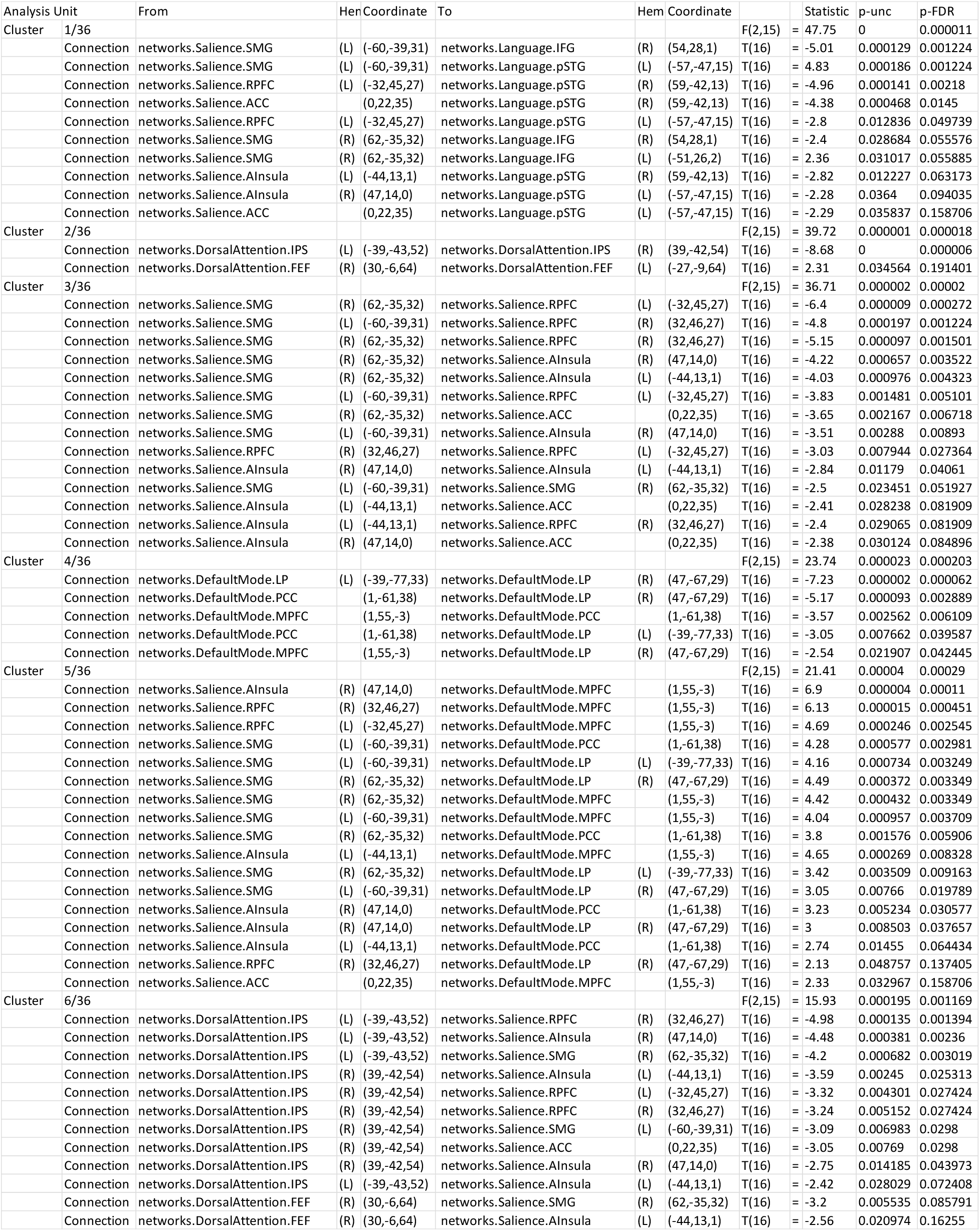
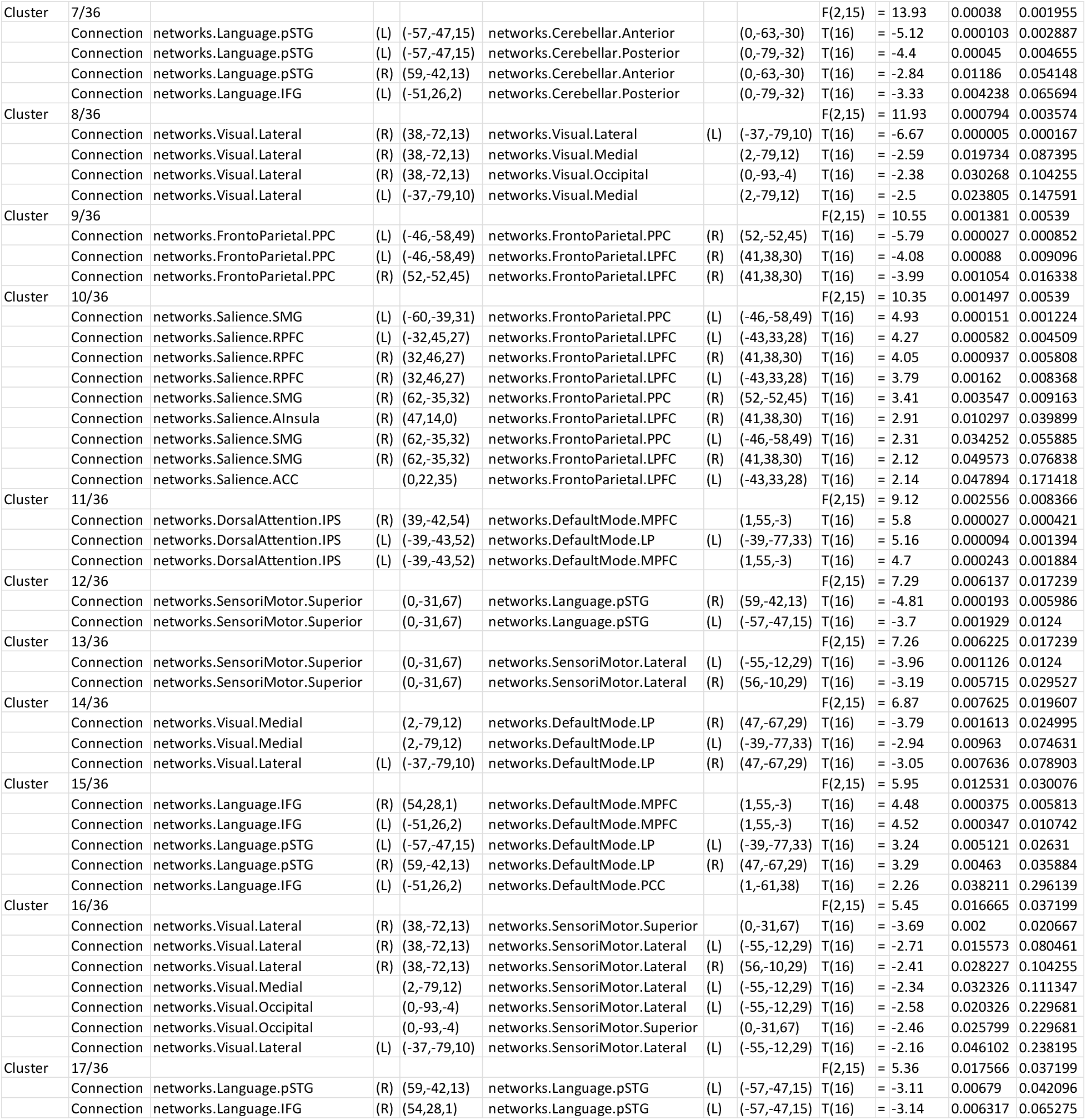
Propofol vs. baseline ROI-to-ROI functional connectivity results.

**Table S6:**
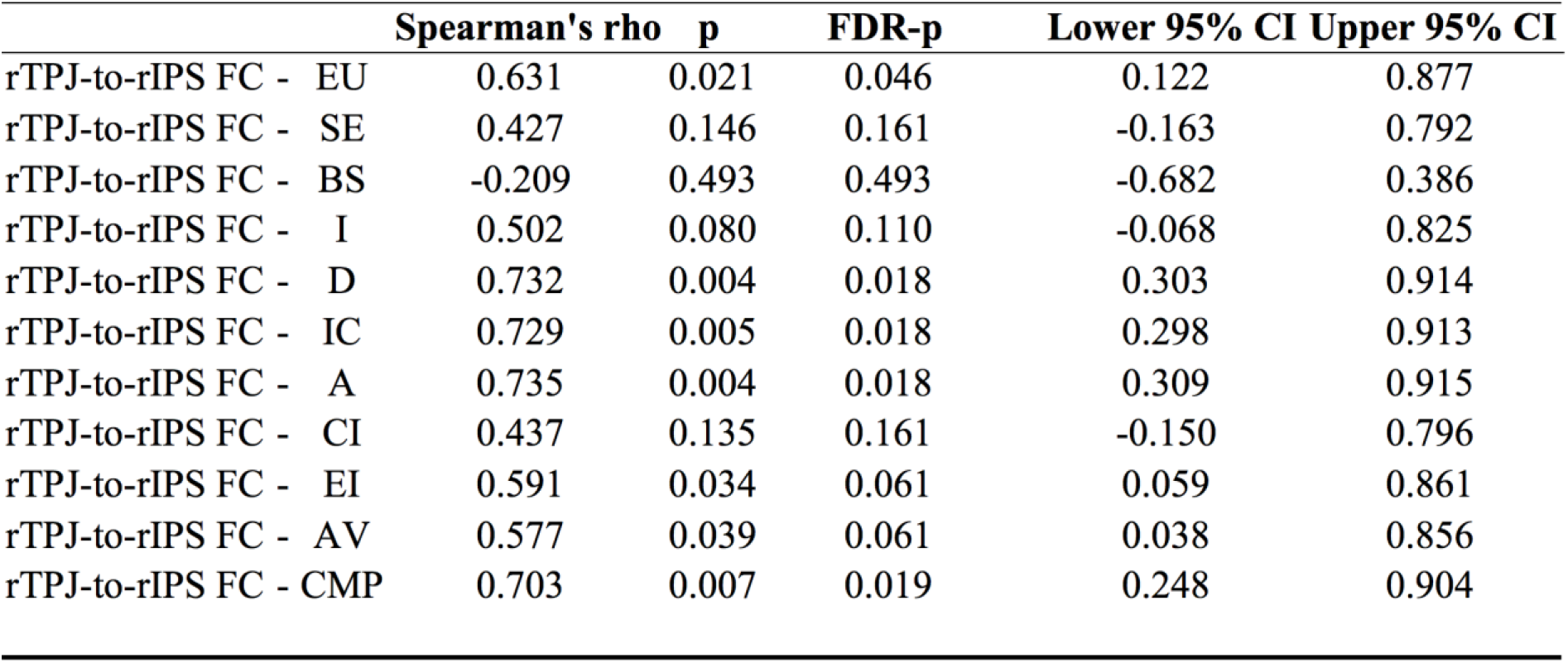
Correlation between right temporoparietal junction-to-right intraparietal sulcus functional connectivity changes and 11D-altered states questionnaire score changes in nitrous oxide data.

|  | **Spearman's rho** | **p** | **FDR-p** | **Lower 95% CI** | **Upper 95% CI** |
| --- | --- | --- | --- | --- | --- |
| rTPJ-to-rIPS FC - EU | 0.631 | 0.021 | 0.046 | 0.122 | 0.877 |
| rTPJ-to-rIPS FC - SE | 0.427 | 0.146 | 0.161 | -0.163 | 0.792 |
| rTPJ-to-rIPS FC - BS | -0.209 | 0.493 | 0.493 | -0.682 | 0.386 |
| rTPJ-to-rIPS FC - I | 0.502 | 0.080 | 0.110 | -0.068 | 0.825 |
| rTPJ-to-rIPS FC - D | 0.732 | 0.004 | 0.018 | 0.303 | 0.914 |
| rTPJ-to-rIPS FC - IC | 0.729 | 0.005 | 0.018 | 0.298 | 0.913 |
| rTPJ-to-rIPS FC - A | 0.735 | 0.004 | 0.018 | 0.309 | 0.915 |
| rTPJ-to-rIPS FC - CI | 0.437 | 0.135 | 0.161 | -0.150 | 0.796 |
| rTPJ-to-rIPS FC - EI | 0.591 | 0.034 | 0.061 | 0.059 | 0.861 |
| rTPJ-to-rIPS FC - AV | 0.577 | 0.039 | 0.061 | 0.038 | 0.856 |
| rTPJ-to-rIPS FC - CMP | 0.703 | 0.007 | 0.019 | 0.248 | 0.904 |

## Notes

### Competing Interest Statement

The authors have declared no competing interest.

## References

1. D. E. Nichols, Psychedelics. Pharmacol. Rev. 68, 264–355 (2016).

2. V. Jevtović-Todorović, S. M. Todorović, S. Mennerick, S. Powell, K. Dikranian, N. Benshoff, C. F. Zorumski, J. W. Olney, Nitrous oxide (laughing gas) is an NMDA antagonist, neuroprotectant and neurotoxin. Nat. Med. 4, 460–3 (1998).

3. R. I. Block, M. M. Ghoneim, V. Kumar, D. Pathak, Psychedelic effects of a subanesthetic concentration of nitrous oxide. Anesth. Prog. 37, 271–6 (1990).

4. W. James, Review of “The Anaesthetic Revelation and the Gist of Philosophy.” Atl. Mon. 33, 627–628 (1874).

5. E. R. John, L. S. Prichep, W. Kox, P. Valdés-Sosa, J. Bosch-Bayard, E. Aubert, M. Tom, F. di Michele, L. D. Gugino, F. DiMichele, Invariant reversible QEEG effects of anesthetics. Conscious. Cogn. 10, 165–83 (2001).

6. B. L. Foster, D. T. J. Liley, Effects of nitrous oxide sedation on resting electroencephalogram topography. Clin. Neurophysiol. 124, 417–23 (2013).

7. K. J. Pavone, O. Akeju, A. L. Sampson, K. Ling, P. L. Purdon, E. N. Brown, Nitrous oxide-induced slow and delta oscillations. Clin. Neurophysiol. 127, 556–564 (2016).

8. J.-H. Ryu, P.-J. Kim, H.-G. Kim, Y.-S. Koo, T. J. Shin, Investigating the effects of nitrous oxide sedation on frontal-parietal interactions. Neurosci. Lett. 651, 9–15 (2017).

9. A. Pelentritou, L. Kuhlmann, J. Cormack, S. Mcguigan, W. Woods, S. Muthukumaraswamy, D. Liley, Source-level Cortical Power Changes for Xenon and Nitrous Oxide-induced Reductions in Consciousness in Healthy Male Volunteers. Anesthesiology. 132, 1017–1033 (2020).

10. X. C. E. Vrijdag, H. van Waart, S. J. Mitchell, J. W. Sleigh, An Electroencephalogram Metric of Temporal Complexity Tracks Psychometric Impairment Caused by Low-dose Nitrous Oxide. Anesthesiology. 134, 202–218 (2021).

11. N. Dashdorj, K. Corrie, A. Napolitano, E. Petersen, R. P. Mahajan, D. P. Auer, Effects of subanesthetic dose of nitrous oxide on cerebral blood flow and metabolism: a multimodal magnetic resonance imaging study in healthy volunteers. Anesthesiology. 118, 577–86 (2013).

12. E. Studerus, A. Gamma, F. X. Vollenweider, Psychometric evaluation of the altered states of consciousness rating scale (OAV). PLoS One. 5, e12412 (2010).

13. M. E. Liechti, P. C. Dolder, Y. Schmid, Alterations of consciousness and mystical-type experiences after acute LSD in humans. Psychopharmacology (Berl). 234, 1499–1510 (2017).

14. S. Arzy, G. Thut, C. Mohr, C. M. Michel, O. Blanke, Neural basis of embodiment: distinct contributions of temporoparietal junction and extrastriate body area. J. Neurosci. 26, 8074–81 (2006).

15. P. E. Vlisides, T. Bel-Bahar, A. Nelson, K. Chilton, E. Smith, E. Janke, V. Tarnal, P. Picton, R. E. Harris, G. A. Mashour, Subanaesthetic ketamine and altered states of consciousness in humans. Br. J. Anaesth. 121, 249–259 (2018).

16. C. Koch, M. Massimini, M. Boly, G. Tononi, Neural correlates of consciousness: progress and problems. Nat. Rev. Neurosci. 17, 307–21 (2016).

17. Z. Huang, X. Liu, G. A. Mashour, A. G. Hudetz, Timescales of Intrinsic BOLD Signal Dynamics and Functional Connectivity in Pharmacologic and Neuropathologic States of Unconsciousness. J. Neurosci. 38, 2304–2317 (2018).

18. P. Nagele, A. Duma, M. Kopec, M. A. Gebara, A. Parsoei, M. Walker, A. Janski, V. N. Panagopoulos, P. Cristancho, J. P. Miller, C. F. Zorumski, C. R. Conway, Nitrous Oxide for Treatment-Resistant Major Depression: A Proof-of-Concept Trial. Biol. Psychiatry. 78, 10–18 (2015).

19. P. Nagele, B. J. Palanca, B. Gott, F. Brown, L. Barnes, T. Nguyen, W. Xiong, N. C. Salloum, G. D. Espejo, C. N. Lessov-Schlaggar, N. Jain, W. W. L. Cheng, H. Komen, B. Yee, J. D. Bolzenius, A. Janski, R. Gibbons, C. F. Zorumski, C. R. Conway, A phase 2 trial of inhaled nitrous oxide for treatment-resistant major depression. Sci. Transl. Med. 13, 1376 (2021).

20. D. De Gregorio, A. Aguilar-Valles, K. H. Preller, B. D. Heifets, M. Hibicke, J. Mitchell, G. Gobbi, Hallucinogens in Mental Health: Preclinical and Clinical Studies on LSD, Psilocybin, MDMA, and Ketamine. J. Neurosci. 41, 891–900 (2021).

21. Z. Huang, J. Zhang, J. Wu, G. A. Mashour, A. G. Hudetz, Temporal circuit of macroscale dynamic brain activity supports human consciousness. Sci. Adv. 6, 1–15 (2020).

22. R. L. Carhart-Harris, S. Muthukumaraswamy, L. Roseman, M. Kaelen, W. Droog, K. Murphy, E. Tagliazucchi, E. E. Schenberg, T. Nest, C. Orban, R. Leech, L. T. Williams, T. M. Williams, M. Bolstridge, B. Sessa, J. McGonigle, M. I. Sereno, D. Nichols, P. J. Hellyer, P. Hobden, J. Evans, K. D. Singh, R. G. Wise, H. V. Curran, A. Feilding, D. J. Nutt, Neural correlates of the LSD experience revealed by multimodal neuroimaging. Proc. Natl. Acad. Sci. U. S. A. 113, 4853–4858 (2016).

23. Z. Huang, J. Zhang, J. Wu, P. Qin, X. Wu, Z. Wang, R. Dai, Y. Li, W. Liang, Y. Mao, Z. Yang, J. Zhang, A. Wolff, G. Northoff, Decoupled temporal variability and signal synchronization of spontaneous brain activity in loss of consciousness: An fMRI study in anesthesia. Neuroimage. 124, 693–703 (2016).

24. Z. Huang, J. Zhang, J. Wu, X. Liu, J. Xu, J. Zhang, P. Qin, R. Dai, Z. Yang, Y. Mao, A. G. Hudetz, G. Northoff, Disrupted neural variability during propofol-induced sedation and unconsciousness. Hum. Brain Mapp. 39, 4533–4544 (2018).

25. S. Whitfield-Gabrieli, A. Nieto-Castanon, Conn: a functional connectivity toolbox for correlated and anticorrelated brain networks. Brain Connect. 2, 125–41 (2012).

26. J. Chumbley, K. Worsley, G. Flandin, K. Friston, Topological FDR for neuroimaging. Neuroimage. 49, 3057–64 (2010).

27. R. B. Mars, J. Sallet, U. Schüffelgen, S. Jbabdi, I. Toni, M. F. S. Rushworth, Connectivity-based subdivisions of the human right “temporoparietal junction area”: evidence for different areas participating in different cortical networks. Cereb. Cortex. 22, 1894–903 (2012).

28. Y. Liu, J. H. Gao, M. Liotti, Y. Pu, P. T. Fox, Temporal dissociation of parallel processing in the human subcortical outputs. Nature. 400, 364–7 (1999).

29. G. Tononi, O. Sporns, G. M. Edelman, A measure for brain complexity: relating functional segregation and integration in the nervous system. Proc. Natl. Acad. Sci. U. S. A. 91, 5033–7 (1994).

30. G. Deshpande, S. LaConte, S. Peltier, X. Hu, Integrated local correlation: A new measure of local coherence in fMRI data. Hum. Brain Mapp. 30, 13–23 (2009).

31. M. J. Jafri, G. D. Pearlson, M. Stevens, V. D. Calhoun, A method for functional network connectivity among spatially independent resting-state components in schizophrenia. Neuroimage. 39, 1666–1681 (2008).

32. K. J. Worsley, S. Marrett, P. Neelin, A. C. Vandal, K. J. Friston, A. C. Evans, A unified statistical approach for determining significant signals in images of cerebral activation. Hum. Brain Mapp. 4, 58–73 (1996).

